# Context-Dependent Metabolic Adaptation in Microbial Communities: From Monocultures to Complex Ecological Interactions

**DOI:** 10.1101/2025.10.22.683057

**Authors:** Anna G. Burrichter, Johannes Zimmermann, Chen Meng, Christina Ludwig, Christoph Kaleta, Bärbel Stecher

## Abstract

Microbial communities perform myriad functions in various environments, including the mammalian gut. These functions are largely based on the metabolic activity of microbial communities, which arises from the individual members’ contributions and their interactions. However, microbial functions and interactions are often inferred from simplified systems such as pure or pairwise cultures. Larger communities, in particular, are mainly analysed by metagenomics and the activity is deduced from the genetic potential of their community members. Although these approaches allow detailed insights into compositions, pairwise interactions and potential functions, they neglect the complexities of microbial communities, as evident in higher-order relationships, competition outcomes and the regulatory influence of environmental factors and other community members. Here, we study a mouse gut community along a gradient of complexity - from monocultures, to in vitro communities, to the mouse gut, to probe how bacteria adapt metabolically to their biotic and abiotic environments. Using metaproteomics to analyze realized bacterial niches and metabolic modeling to infer community interactions, we found that bacteria substantially modify their carbon source usage, biosynthetic activities, and protein allocation when grown in communities versus isolation. Furthermore, communities themselves adapted to their growth environment, changing in functionality as well as in the metabolic network between members. Host-associated communities isolated from the mouse gut invested in resource acquisition and utilized a broader spectrum of carbon sources, resulting in a large increase of predicted interactions between them. Our findings demonstrate the remarkable adaptability of microbes and microbial communities across environmental contexts. This underscores the critical need to study microbiomes under near-natural conditions and incorporate context-specific data into functional analyses.

## Introduction

Microorganisms are fundamental to nearly every ecosystem on earth^1^ and exert profound influences when associated with hosts, such as humans, where gut and skin microbes are indispensable for maintaining health^2,3^. The functioning of microbes, however, can only be fully understood by examining them as communities, where collective properties—such as metabolite and vitamin production, colonization resistance against pathogens, and complex substrate degradation—emerge from intricate networks formed through species interactions^4,5^.

While community composition is increasingly accessible through sequencing technologies, understanding microbial community functioning presents two significant challenges. First, composition alone rarely translates directly into function, even with multi-omics data available^6^. This is due to emergent properties caused by higher-order interactions. For instance, certain metabolic capabilities may only emerge when specific combinations of species interact, revealing new molecules that enable bacteria to adapt to complex environments. Second, microbial activities vary substantially across environmental conditions and ecological contexts^7–13^, limiting the applicability of insights gained from single-species studies or simplified co-cultures^5,14^. Addressing these challenges requires novel approaches that examine communities across diverse conditions. For example, combining theoretical models with well-characterized, simplified microbial communities offers a practical approach to unraveling complex ecological interaction^15,16^.

The Oligo-Mouse-Microbiota (OMM^12^) is a defined community widely used for research into gut physiology ^12,17–19^, colonisation resistance^20,21^ and immune system development^22–24^. This defined bacterial community consists of 12 genome-sequenced strains representative of the gut microbiome^25^ of mice. The OMM^12^ confers colonisation resistance to human pathogenic *Salmonella enterica* serovar Typhimurium infection based on resource competition^20^.

Analysis of its internal metabolic network has begun using in vitro pairwise competition assays, dropout communities and metabolomic profiling of single members^7,26,27^: Weiss et al. tested the growth of single members in each other’s spent media, systematically mapping for each pair how species hinder or enhance each other’s growth due to effects such as pH modification, niche overlap or toxin production^26^. Pérez Escriva et al. analysed the metabolome of each species and inferred a putative crossfeeding network from these data^27^. In a later study, Weiss et al. then profiled community dynamics by analysing drop-out communities (OMM^11^ communities missing one species) across different culturing media and in gnotobiotic mice to identify keystone species whose absence has pronounced effects on the abundance of other members or the production of metabolites. Here, the authors found that these effects can vary drastically between different environments, raising the idea that community function is context-dependent^7,28,29^. While these approaches have advanced our understanding of the individual species and their pairwise and community-level interactions, they fall short in capturing the complex processes and metabolic adaptations in individual community members arising from higher-order interactions and the realized niche occupancy within a community context in different environmental settings.

Systems biology approaches such as genome-based metabolic modelling of single bacteria and communities can predict microbial metabolism and community structure^30,31^ and resolve the molecular basis of interactions that are not directly accessible through experimental measurements^32,33^. These models are based on the overall genomic potential of organisms and predict metabolic activity based on mathematical optimization, thus capturing the fundamental and not necessarily the realized metabolic activity. They also allow to integrate and interpret large meta-omic datasets^34,35^, since additional data such as metabolomic^36^, metatranscriptomic^36–38^ or metaproteomic^39,40^ measurements can constrain possible solutions, thereby enhancing their predictive accuracy (reviewed comprehensively by Heinken et al.^41^). Integrative approaches have been used successfully to predict disease-based changes of community metabolism or short-chain fatty acid production profiles of individual gut microbiomes^42–46^, detailed cross-feeding mechanisms between bacterial species^47,48^, or to identify specific prebiotics^49^.

Yet, the gap between microbial physiology at the cellular level and the behaviour of microbiomes as an ecosystem is rarely bridged^50^, hindering the full understanding of how environmental changes impact microbial communities. Here, we present an integrative systems biology approach to elucidate community interaction and functions of OMM^12^, based on metabolic changes of individual member species. We generated comprehensive metaproteomic datasets of OMM^12^ in different growth environments of varying complexity and integrated these data with genome-scale metabolic models to simulate community interactions and metabolic activities. We observe a significant reconfiguration of species-specific metabolic activities and a marked increase in interspecies interactions within host-associated communities compared to in vitro culture conditions. This is accompanied by distinct metabolic reprogramming in the intestinal environment, where microbial communities prioritize resource acquisition over growth maximization.

## Results

### OMM^12^ community members adapt protein expression in response to the microbial environment

First, we investigated the adaptation of the individual OMM^12^ bacteria to the microbial community environment. To this end, we compared proteomic data of single OMM^12^ strains grown in AF medium to metaproteomics of the OMM^12^ community grown under the same conditions (**Fig. 1A**). AF medium is a complex anaerobic culture medium that was previously identified to best support growth of individual OMM^12^ strains^7^. Individual strains were cultured in AF media to mid-log phase (n=3) (**Table S1**) and stabilised OMM^12^ communities for 24 hours. Samples were prepared and analysed as described in the Methods section. Mass spectra were searched using MaxQuant against a peptide library generated from genomes of all 12 OMM^12^ members and protein signals were quantified using the label-free quantification (LFQ) algorithm. Between 987 (*Limosilactobacillus reuteri* I49) and 1909 (*Enterocloster clostridioformis* YL32) proteins were detected per species in their respective monocultures (**Fig. 1B**). In the community, between 869 and 168 proteins were detected per species, with four additional species being so low abundant that less than 20 proteins could be recovered for each of them (*Bifidobacterium animalis* YL2, *Acutalibacter muris* KB18, *Akkermansia muciniphila* YL44 and *L. reuteri* I49; these species are excluded from the species-resolved analysis below) (**Fig. 1B** and **Fig. S1** for the community composition analysed by qPCR). Since these datasets were highly uneven in depth for each species, we analysed proteins based on their rank in each species’ total proteome.

**Figure 1:**
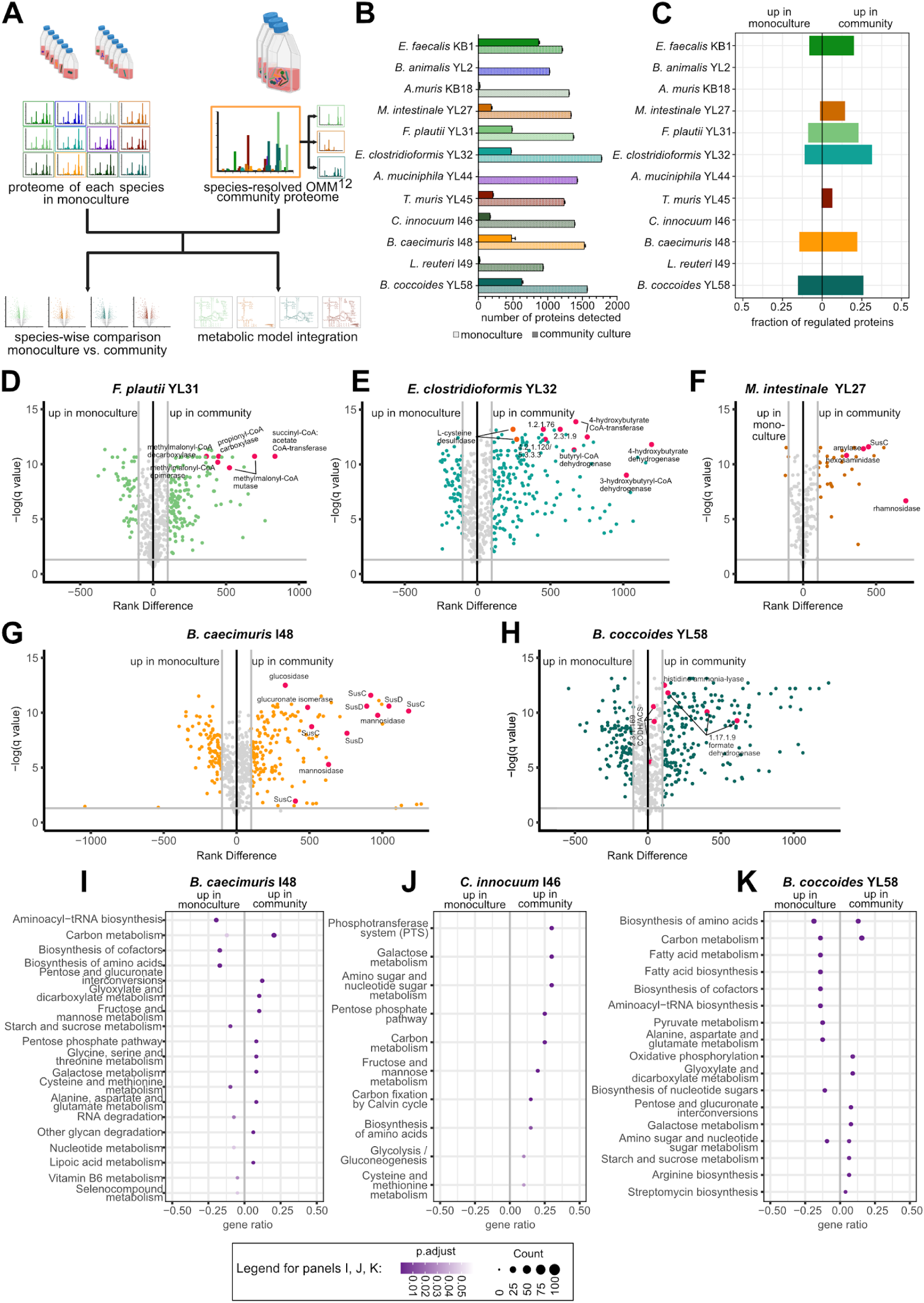
**Bacterial metabolism is influenced by the biotic environment.** A: Systematic overview of the experimental approach for the comparison of single vs. community culture of each species. B: Number of proteins (mean +/- SD) detected for each species in monoculture (individual strain colours) and in the community (pink). C: Fraction of proteins (of all proteins detected for the respective species in community culture) that showed a rank change of 100 or more when ranking proteins by mean MS intensity (LFQ). Proteins to the left of center rank higher in the monoculture, proteins to the right rank higher in the community. D-H: Volcano plot of proteins of individual species in monoculture vs. community culture: x axis: rank difference monoculture - community culture; y axis: q value of unpaired multiple t-tests with a false discovery rate correction for multiple comparisons (FDR cutoff 1% with the two-stage step-up method of Benjamini, Krieger and Yekutieli^51^); grey lines mark a rank change of 100 and a q value of 0.05. Full annotations of marked proteins can be found in Table S2. Multiple data points with the same description account for different protein subunits or homologs. D: Volcano plot for *Flavonifractor plautii* YL31, enzymes of propionate formation are marked in red. E: Volcano plot for *Enterocloster clostridioformis* YL32, enzymes of butyrate formation are marked in red and enzymes of sulfide formation in orange (2.3.1.9: acetyl-CoA C-acetyltransferase; 4.2.1.120 / 5.3.3.3: 4-hydroxybutyryl-CoA dehydratase / vinylacetyl-CoA-Delta-isomerase; 1.2.1.76: succinate-semialdehyde dehydrogenase). F: Volcano plot for *Muribaculum intestinale* YL27, enzymes for polysaccharide utilisation are marked in red. G: Volcano plot for *Bacteroides caecimuris* I48, enzymes for polysaccharide utilisation are marked in red. H: Volcano plot for *Blautia coccoides* YL58, enzymes of the Wood-Ljungdahl pathway and histidine ammonia-lyase are marked in red (CODH/ACS: carbon monoxide dehydrogenase/acetyl-CoA synthase complex). I: Pathway enrichment (KEGG pathways) of significantly regulated proteins with a rank change >100 for *B. caecimuris* I48. J: Pathway enrichment (KEGG pathways) of significantly regulated proteins with a rank change >100 for *Clostridium innocuum* I46. I: Pathway enrichment (KEGG pathways) of significantly regulated proteins with a rank change >100 for *B. coccoides* YL58. Full enrichment data can be found in Supplementary Table S3.

Overall, more than 1000 proteins showed a rank change of 100 or more, suggesting regulation by the biotic context (**Fig. 1C**). We observed higher expression of core metabolic pathways in multiple members of the OMM^12^ when grown as a community, with the enzymes of these pathways ranking significantly higher within the proteome under community cultivation conditions. For example, *Flavonifractor plautii* YL31 and *E. clostridioformis* YL32 upregulated propionate and butyrate production, respectively (**Fig. 1D, E**). On the substrate side, Bacteroidales species *Bacteroides caecimuris* I48 (**Fig. 1F**) and *Muribaculum intestinale* YL27 (**Fig. 1G**) expressed various polysaccharide-utilizing enzymes more highly in the community context. In contrast, *Clostridium innocuum* I46 expressed phosphotransferase systems for more specific carbon sources and *Blautia coccoides* YL58 increased expression of the Wood-Ljungdahl pathway (**Fig. 2H**). Classifying these proteins into KEGG pathways (**Fig. S3**), the most common group of regulated proteins were part of carbon and amino acid metabolism, closely followed by biosynthesis of amino acids and cofactors. Across all species, these functions were both up- and downregulated (**Fig. S3**).

**Figure 2:**
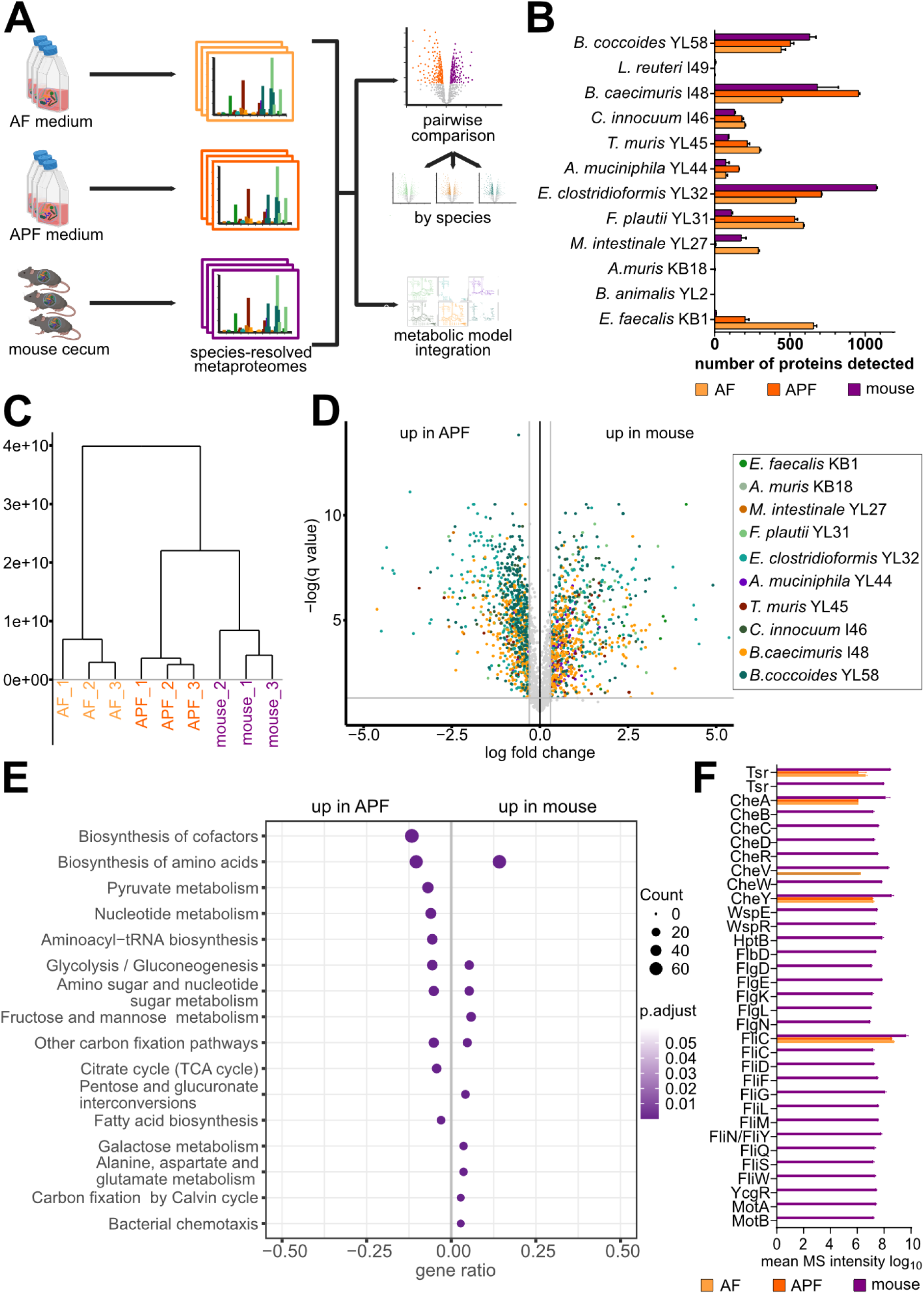
**Bacterial metabolism is influenced by the abiotic environment.** A: Systematic overview of the experimental approach for the comparison of OMM^12^ in vitro and in the murine gut. B: number of proteins (mean +/- SD) detected for each species in each sample. C: Hierarchical clustering analysis of community proteomes using Euclidean distances and complete linkage clustering. D: Volcano plot of differentially regulated proteins between OMM12 cultured in vitro in APF medium and extracted from the mouse cecum. x axis: log fold change of species-normalised protein intensities; y axis: q value of unpaired multiple t-tests with a false discovery rate correction for multiple comparisons (FDR cutoff 5% ; grey lines mark a fold change of 2 and a q value of 0.05. E: KEGG enrichment of differentially expressed proteins (fold change > 2) between APF in vitro medium and the mouse cecum for all species (AF comparisons and enrichment by species in Supplementary Fig. S9 and S10, enrichment data in Supplementary Table S4) For clarity, only the top 10 enriched pathways in each direction are shown and the pathways “carbon metabolism” and “ribosome” have been removed. F: expression levels under all three conditions of the operons for chemotaxis and flagellar machinery in *Enterocloster clostridioformis* YL32 (mean MS intensity +/- SD). Full gene annotations in Table S5.

This effect could be resolved by enriching differentially regulated enzymes by species (**Fig. 1I-K** and **Fig. S4)**. Biosynthesis of cellular components (amino acids and cofactors) was downregulated in the community context for some members such as *B. caecimuris* I48 (**Fig. 1I**) but upregulated for others like *Enterococcus faecalis* KB1 (**Fig. S4A**) and *C. innocuum* I46 (**Fig. 1J**). Alanine, aspartate and glutamate metabolism was downregulated by *B. coccoides* YL58 (**Fig. 1K**) and upregulated by *B. caecimuris* I48 (**Fig. 1I**) in the community. Utilization pathways for specific carbon sources such as amino sugars, galactose, sucrose or pentoses and glucuronates were among the proteins upregulated in the community as the number of available carbon sources narrows for each individual species due to potential substrate competition leading to niche switching from glucose to other carbon sources.

Comparison of signal peptides of carbohydrate active enzymes (CAZymes) revealed further differences between mono and community proteins, indicating a possible increased importance of extracellular enzymes in *B. caecimuris* I48 and *M. intestinale* YL27 under community conditions (**Fig. S5**). In conclusion, we found that upon integration into a microbial community, individual bacterial species rewire their metabolism; and pathways for biosynthesis, amino acid metabolism and carbon source utilisation are differentially regulated between species. This possibly reflects increased competition for accessible nutrients or increased crossfeeding^26^.

### OMM^12^ community members adapt protein expression in response to the environment of the murine gut

Next, we investigated the adaptation of OMM^12^ communities to different culturing conditions. The culture medium has previously been shown to significantly influence community structure and identity of keystone members^7^. Although in vitro models are valuable for dissecting mechanistic aspects of in vivo phenotypes, their simplified environments often fail to replicate the complex host factors and microenvironments present in living organisms, leading to differences in bacterial ecology and limited translational potential. We compared the established standard rich anaerobic culture medium for OMM^12^ (AF) (n=3), a modified version developed to more closely resemble the community composition in the mouse gut^26^ (APF; polysaccharide-rich) (n=3) and cecum samples of mice stably colonized with OMM^12^ (n=3) (experimental overview see **Fig. 2A**, for species composition of these communities analyzed by qPCR, see **Fig. S6**). Protein counts for each species are depicted in **Fig. 2B** - again, species *B. animalis* YL2, *A. muris* KB18 and *L. reuteri* I49 could not be detected beyond single proteins and were thus excluded from further analysis. Indeed, the proteome of the OMM^12^ community grown in APF was closer to the metaproteome from the mouse cecum in a hierarchical clustering analysis (**Fig. 2C**). This could partially be explained by the expression of enzymes targeted at utilization of additional carbon sources such as *B. caecimuris* I48 xylanases and inulinase induced by the availability of these complex carbohydrates in APF (**Fig. S7A**). Still, significant differences in protein abundance patterns between conditions remained (**Fig. 2D**, comparisons for AF vs. APF and AF vs. mouse in **Fig. S8**). Enrichment analysis of differentially expressed proteins over all species (**Fig. 2G**, species-resolved enrichments in **Fig. S10**) showed that metabolic pathways for a wider variety of carbon sources (pentoses, amino sugars, galactose) are more highly expressed in the cecum while culturing in APF enriched for functions of central carbon metabolism such as glycolysis and the TCA cycle. Whole proteins are not as abundant in liquid media as compared to the gut as they are often removed during media processing. Therefore, for example peptidases of *C. innocuum* I46 may be found only in vivo while degradation pathways for single amino acids occur under all conditions (**Fig. S7B, C**). Other carbon sources that might be missing in vitro as indicated by expressed pathways are glucuronates and pentoses (**Fig. 2E**) as well as hydrophobic cell components such as glyceroplipids and lipoic acid (**Fig. S10C, H**). A further function expressed only in vivo was chemotaxis and motility, which likely reflects the lack of spatial organisation and surfaces in well-mixed liquid media (**Fig. 2E** and **S10D**). Specifically, flagella genes of *E. clostridioformis* YL32 were found to be upregulated in the mouse cecum (**Fig. 2F**). Summing up this part, while modifying the growth medium can align protein expression and bacterial function more closely with in vivo conditions, discrepancies persist due to differences in available carbon sources and structural environmental factors.

### The host-associated community follows a resource acquisition strategy

Cellular resources spent on protein synthesis could be spent in several, potentially opposing, ways. It is thus interesting to ask how much specific groups of functionally related proteins contribute to the total protein mass expressed in a given bacterium at a given time point. The protein mass fraction can be estimated from quantitative proteomic data by summing the measured mass spectrometric intensities of all proteins involved in a given functional group compared to the total measured intensity. Building on this, we analysed the allocation of proteomic resources to different functions to elucidate microbial strategies under various growth conditions. For the comparison of monoculture and community growth across species (**Fig. 3A**), we found a similar picture as for the pathway enrichment in **Fig. S3**: Growth in a community setting led to relatively higher protein investment in energy production (COG category **C**), carbon source acquisition (**G**), and translation (**J**), while transcription (**K**), replication (**L**) were reduced. The COG categories **E**, **F**, and **H**, which contain biosynthetic functions, were differentially regulated between species (**Fig. S11**).

**Figure 3:**
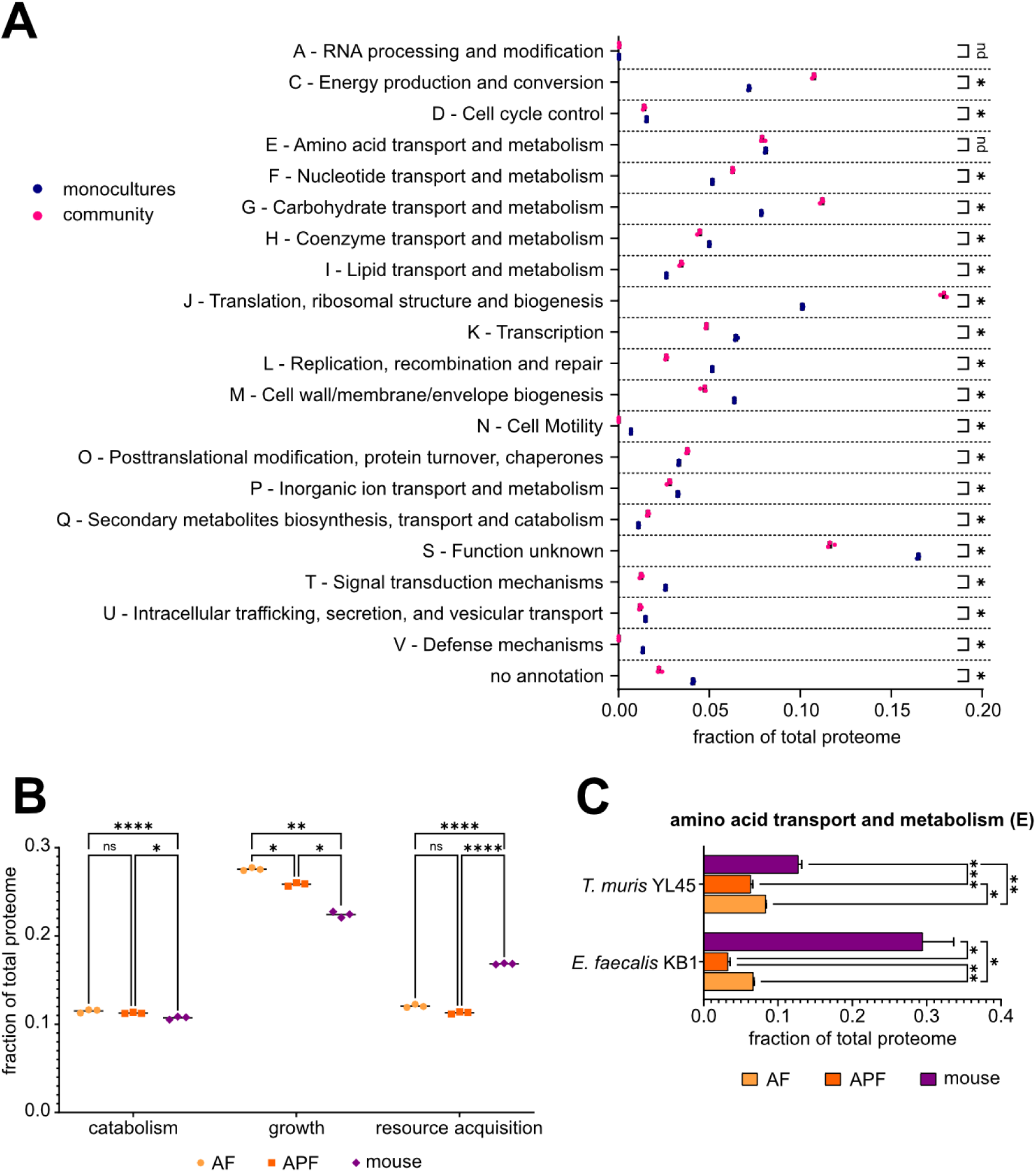
Proteome allocation under different growth conditions. A: proteome fraction per COG category in monoculture (blue) and in vitro community culture (pink) across all species (single species see Fig. S11). B: proteome category groups for AF and APF in vitro cultivations and mouse cecum over all species. COG categories per group: catabolism: C; growth: D, J, K, L; resource acquisition: G, N, T. (for data for each species see Fig. S12, single categories see Fig. S13). C: expression levels of COG category E: amino acid transport and metabolism for AF and APF in vitro cultivations and mouse cecum for *Turicimonas muris* YL45 and *Enterococcus faecalis* KB1. Statistics: A: multiple unpaired t-tests with a false discovery rate correction for multiple comparisons (FDR cutoff 0.5% with the two-stage step-up method of Benjamini, Krieger and Yekutieli^51^) B,C:Two-way ANOVA followed by Tukey’s test with confidence interval 95%.

The analysis of how the microbial community allocated its proteomes when grown in vitro and in vivo conditions revealed a distinct strategy employed by the community in a host-associated environment. The host-associated microbiome demonstrated a clear shift toward utilization of resources at the expense of growth (**Fig. 3B**). This was evidenced by a significant reduction in growth-related proteins (COG categories **D**: Cell cycle control, cell division, chromosome partitioning; **J**: Translation, ribosomal structure and biogenesis; **K** Transcription; **L**: Replication, recombination and repair) and concurrent increases in carbohydrate metabolism and transport, signalling and motility proteins (categories **G**, **T**, and **N**) in mouse-associated communities compared to AF and APF in vitro cultivations. The greater use of carbon metabolism, motility, and signaling is, in general, characteristic of a resource acquisition strategy^52^. We therefore propose a trade-off between growth and resource acquisition, which was further supported by the decreased abundance of proteins assigned to energy production and conversion (**C**) in mouse-associated communities (**Fig. 3B**). When considering individual members of the community, some species showed an opposing trend of increased allocation of proteome towards growth and less resource acquisition, evident for *E. faecalis* KB1, *F. plautii* YL31 and *Turicimonas muris* YL45 (**Fig. S12A, C, F**). However, for *E. faecalis* KB1 and *T. muris* YL45, this could be explained by an increased amino acid metabolism (higher category **E**, **Fig. 3C**), as *E. faecalis* KB1 consumes serine and arginine, and *T. muris* YL45 utilizes aspartate and asparagine^7,26^. This aligns with a broader definition of resource acquisition, which also encompasses amino acid metabolism, in addition to carbohydrate metabolism.

### Metabolic models reflect context-specific metabolism

To gain a deeper insight into the metabolic functioning and adaptation of OMM^12^ strains, we utilized curated genome-based metabolic models constrained by proteomic data. We reconstructed metabolic models from genomes using gapseq^53^ and curated pathways involved in fermentation, polysaccharide, and amino acid metabolism. We further considered those proteins as active that showed a median value greater than zero across the three replicates to derive a context-specific model for each organism of the OMM^12^ community using the fastcore algorithm (see Methods). These context-specific models enabled us to observe community adaptation to biotic and abiotic environmental changes, as well as to model the overall community. Analysis of proteome-constrained metabolic models demonstrated that monoculture proteomes had the highest percentage of model genes linked to expressed proteins, while in vivo cultivation showed the lowest, reflecting a reduced proteome resolution in community and host settings (**Fig. S14A**). Interestingly, the proportion of proteins linked to model genes per species remained stable across all conditions (∼25% coverage; **Fig. S14A**). This is similar to the average coverage of genes of a genome in metabolic models, which ranges between 9% and 21% for the OMM^12^ community (**Fig. S14C**), suggesting that metabolism-relevant proteins maintain a consistent representation regardless of proteome resolution. The percentage of proteins assigned to metabolism-related COG categories was about 50% for communities (**Fig. S14B**, red shades). In addition, the number of reactions in metabolic networks varied across conditions, with most reactions in models from monocultures and a decreasing number of reactions in models from vitro and vivo conditions, again reflecting the lower proteome resolution in community settings (**Fig. S15**). We next compared a) the dissimilarity of our models from mono, in vitro and in vivo cultures (within conditions), and b) the dissimilarity of models between those conditions. Within conditions, we found the lowest dissimilarity between species-specific models in monoculture conditions (mean Jaccard dissimilarity 0.6). The highest dissimilarity was found under in vivo conditions (mean Jaccard similarity 0.9) (**Fig. 4A**). When comparing between conditions, the lowest distance was found between in vitro and in vivo models. In contrast, monoculture models were more distant from in vitro and in vivo models, indicating a general increase in model dissimilarity, starting with monoculture and progressing through in vitro to in vivo models. (**Fig. 4B, Fig. S16**). Reaction-based clustering of context-specific metabolic models aligned with growth conditions. Logistic PCA (**Fig. 4C**) revealed the most pronounced differences in monoculture models compared to other conditions, indicating altered metabolic reaction availability in communities. However, species-specific variations were observed for *E. faecalis* KB1 (AF is closer to monoculture), *F. plautii* YL31 (AF and APF are closer to monoculture), and *B. caecimuris* I48 (APF is closer to monoculture) (**Fig. 4B, C**). Species-specific variations were observed for *E. faecalis* KB1 and *F. plautii* YL31, which showed greater changes in APF and in vivo conditions.

**Figure 4:**
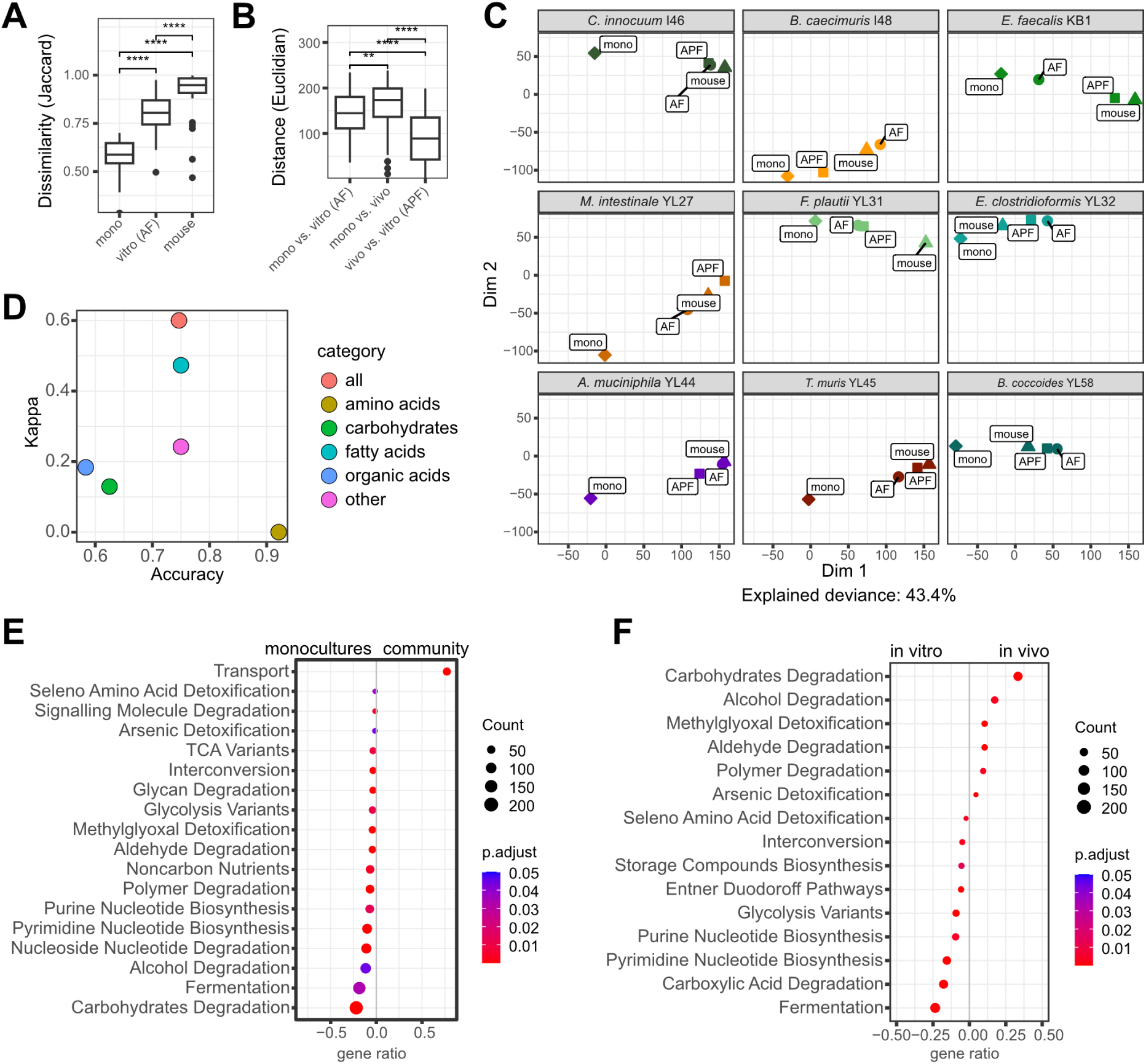
Integration of proteomic data into metabolic models. A: Jaccard similarity of models based on reaction presence within conditions (Wilcoxon rank sum test). B: Euclidian distance of models between conditions based on Logistic PCA coordinates (Wilcoxon rank sum test). C: Logistic PCA of reaction presence for each model under various conditions. D: Validation by comparing model predictions with experimental measurements^7,26,54,55^ (all: overall accuracy). E: Overrepresentation analysis of significant pathways (MetaCyc) for monoculture vs. *in vitro* community (AF medium). F: Overrepresentation analysis of significant pathways for *in vitro* community (APF medium) vs. *in vivo* community.

Validation against previously described^7,25–27,54^ metabolite production and consumption patterns of the individual OMM^12^ members demonstrated >70% prediction accuracy, with particularly high conformity for fatty acids, carbohydrates, and amino acids (**Fig. 4D, Table S6**). Pathway enrichment analysis identified numerous lower abundant pathways when comparing monoculture vs. in vitro community and in vitro community vs. in vivo conditions (**Fig. S14A, Fig. 4E**, **Fig. 4F**). Pathways enriched in the in vitro community compared to monocultures and in vivo compared to in vitro are primarily involved in transport and carbohydrate degradation functions (**Fig. 4E**, **Fig. 4F**). This underscores the concept that both increased resource limitation in the community and host gut environment and host-associated growth drive bacteria to allocate more resources toward acquiring carbon and energy.

### Adaptation leads to changes in bacterial interaction and community ecology

With context-specific models reflecting cells growing in a community setting, we set out to model the metabolic network of the whole community. Therefore, we used context-specific models for each of the three community environments (AF, APF, and the mouse cecum) and simulated their growth as a consortium using BacArena^56^. The starting numbers of each species were set according to qPCR relative abundances (**Fig. S6**) and the resulting growth is depicted in **Fig. 5A** (growth curves in **Fig. S17**). Short-chain fatty acid production patterns agreed with experimentally determined data^7,28,29^ (**Fig. S19, Table S6**). The uptake and release of metabolites is calculated for each simulated cell, allowing the construction of metabolic networks that predict cross-feeding interactions (**Fig. 5B**, **Table S7**). We observed a significantly more complex web of interactions in the in vivo setting compared to both in vitro settings (**Fig. 5C**). Small molecules such as serine, propanal and pyruvate were exchanged solely in vivo, along with polysaccharide degradation products such as melibiose.

**Figure 5:**
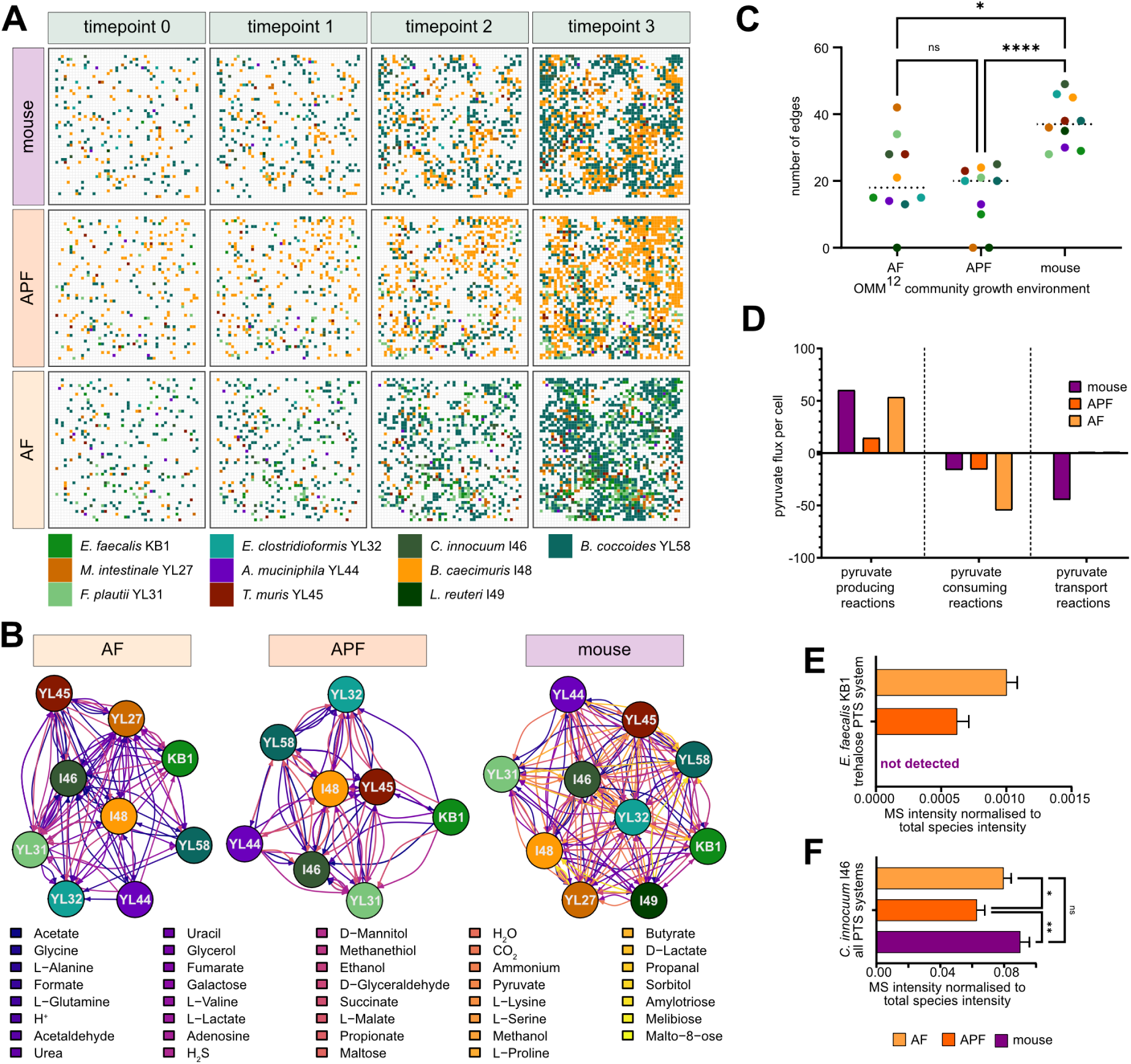
Community simulation using context-specific metabolic models. A: Growth of cells in simulations modelling AF, APF and in vivo growth over three timesteps. The 250x250 grid represents the arena used in the simulation with each grid cell either empty or filled with an organism. B: predicted metabolite exchange networks under all conditions (data in Table S7). C: Connectivity of the networks in B, one-way ANOVA followed by Tukey’s test for multiple comparisons. ****: adj. p < 0.0001. D: pyruvate fluxes in the *Clostridium innocuum* I46 context-specific models under all three community growth conditions (data in Table S8). E: expression levels (mean +/- SD) of the *Enterococcus faecalis* KB1 trehalose-specific phosphotransferase system under all conditions (protein MS intensity normalised to the sum of *E. faecalis* KB1 protein intensities under that specific condition; no protein was detected in the samples from the mouse cecum). F: sum of expression levels (mean +/- SD) of all detected *C. innocuum* I46 phosphotransferase systems under all conditions (protein MS intensity normalised to the sum of *C. innocuum* I46 protein intensities under that specific condition; no trehalose-specific system was detected). Statistics: One-way ANOVA followed by Tukey’s multiple comparisons test; *: p<= 0.05; **: p<= 0.01. Full annotations for the proteins depicted in E and F can be found in Table S9.

As a prominent example, we more closely analysed pyruvate exchange which only appeared in the in vivo simulation using metabolic modelling. Pyruvate as an intermediate of central carbon metabolism is likely to be tightly regulated by carbon source availability. In the mouse gut, predicted pyruvate flux was driven by pyruvate overproduction by *C. innocuum* I46, with the resulting pyruvate taken up by various other members (**Table S8**). By examining all reactions that consume, produce or transport pyruvate in *C. innocuum* I46, we found that intracellular pyruvate production and consumption were uniquely unbalanced in the mouse context-specific model (**Fig. 5C, Table S8**), leading to pyruvate export from the cell.

Intriguingly, we found intracellular pyruvate production and consumption balanced under all conditions. The additional pyruvate produced under in vivo conditions was predicted to be generated as a side product of phosphotransferase systems (PTS), with the highest predicted flux for a trehalose PTS. We therefore hypothesised that under in vivo conditions, the substrate spectrum of *C. innocuum* I46 changed to substrates that are transported via PTS rather than other transport systems. From proteomic data, we could confirm that the trehalose-specific PTS component of *E. faecalis* KB1 is found only under in vitro conditions (AF and APF, **Fig. 5E, Table S9**), potentially leaving this niche open in vivo for *C. innocuum* I46, which expresses significantly more PTS system components in the mouse cecum as compared to APF (**Fig. 5F, Table S9**). This example shows the value of context-specific models to predict the cascading effects of adaptation and metabolic reprogramming. The models can be used in follow-up studies aimed at rewiring community metabolic output by altered nutrient supplies.

## Discussion

Microbial communities exhibit context-dependency of gene expression, ecological relationships and community functions^8–11,29,57–59^, including the OMM^12^ model community studied here^12,26^. We expanded these studies using metaproteomics and mathematical simulation to identify bacterial functions and metabolic pathways that are regulated in response to a complex microbial and host environment. We reveal how focal bacteria dynamically adjust their metabolic activities in response to changing biotic and abiotic conditions, resulting in significant variations in metabolic networks, even when community members remain constant. The microbial community context has a pronounced influence on the proteome of individual bacteria leading to upregulation of carbon-source specific degradation pathways and downregulation of central carbon metabolism as well as species-specific differential regulation of biosynthetic pathways. The host environment triggers the expression of motility systems and provides further carbon sources, again leading to expression of specific degradation pathways. Proteomics constrained metabolic modeling pointed at changes of metabolic interactions with increase in interactions in the murine gut.

Uncovering the mechanisms by which intestinal microbial interaction networks dynamically change across different microbial and nutritional environments enhances our understanding of the molecular basis underlying shifts in the intestinal metaproteome and metabolome, and reveals potential entry points for microbiome engineering to promote intestinal health.

Metaproteomic analysis revealed a substantial redistribution of metabolic activities between species when comparing bacteria grown in monoculture to those in communities across different in vitro cultivation media and the mouse cecum. Bacteria grown in community settings demonstrated marked adaptation in their carbon and energy source utilization profiles, as well as their biosynthetic activities, possibly due to more limited resources while growing in communities. In line with this, the sizes of context-specific metabolic networks decreased from mono to vivo conditions (**Fig. S14A**), which further suggests a metabolic specialization. This increased specialization during the transition from monoculture to in vitro communities has also been shown in two-member co-cultures of gut bacteria^29,57,59–62^. Kamrad et al. studied 104 pairs of human gut bacteria compared to the respective monocultures by proteomics and found that among the globally responsive proteins, i.e. the proteins that are upregulated as a result of coculture across all tested pairs, are enriched in KEGG pathways for carbohydrate metabolism. The same pattern was found in our study, with carbon metabolism upregulated across all species when comparing monocultures to community culture. The metabolism of specific carbon sources and amino acids was also regulated differentially between species (**Fig. 1I-K** and **Fig. S3**). We could also observe the distribution of biosynthetic activities, e.g. for amino acids and cofactors among community members (**Fig. 1I-K** and **Fig. S4**), a behaviour that has been noted for a methanogenic community^63^ and pairs of gut bacteria^57^ and may be due to proteome efficiency^64^. In addition, we identified several predicted extracellular proteins such as glycosidases, esterases and peptidases, that were more abundant in communities, which has also been reported for the human gut microbiome^65^. These findings underscore the need of caution when extrapolating phenotypic data from monocultures to infer bacterial phenotypes in more complex or even host-associated communities.

Notably, when comparing communities between different in vitro cultivation media and the mouse cecum, metabolic activities were again redistributed between species, as has been shown before in changing nutritional environments^12,28,66^. Between glucose-rich and polysaccharide-rich in vitro medium, bacteria adapted to the change in carbon sources according to their genetic potential. In vivo, further complex carbon sources were utilised and protein investment into central carbon metabolism was reduced. Furthermore, proteins relevant for cell motility were only expressed in vivo, reflecting the more structured and less mixed environment.

Our integrative proteome allocation approach provided a global perspective on protein resource utilization within microbial communities, enabling the identification of ecological strategies based on functional analysis of proteome sectors. This analysis revealed that the proteomes of in vitro communities are characterized by growth-oriented processes. In contrast, carbohydrate metabolism, transport mechanisms, signaling and motility processes become more critical under in vivo conditions. In microbial life history frameworks, chemotaxis, which is linked to signaling, and motility are typical foraging traits^67,68^. Foraging is conceptualized to play a vital role in conditions of spatial and temporal variability^67^. In organisms with flagellum motility, the COG categories “signal transduction mechanisms” (linked to chemotaxis) and “carbohydrate transport and metabolism” were consistently found to be enriched; a co-occurrence of traits that the authors summarize as a resource acquisition strategy^52^. In our study, we also identified these traits as co-occurring and classified them accordingly. Additional support for such a resource acquisition life history strategy comes from a global analysis, which described the combined presence of motility, carbohydrate metabolism, and larger genome traits in soil bacteria employing a competitor with high environmental responsiveness strategy^69^. Communities with these attributes were associated with soils from wet and stable climates, environmental factors that are also typical of the gut environment. In addition, bacterial motility and chemotaxis genes have been described as prevalent in the photic zone of the ocean, which exhibits steep gradients of light, temperature, macronutrient, and trace-metal concentrations^70^. Similarly, the mammalian gastrointestinal tract is an environment with a rich biogeography shaped by gradients of pH, oxygen, peristalsis, antimicrobial peptides, mucus, and nutrients^71^. We therefore hypothesize that bacterial adaptation towards resource acquisition under in vivo conditions is a consequence of the biogeographical diversity and nutrient gradients characteristic of host-associated microbiomes.

We further integrated the proteomic data into metabolic models to generate context-specific models and simulate community growth under the varying conditions. The application of metabolic modeling proved valuable for simulating microbial community growth under varying conditions and uncovering mechanisms underlying community dynamics. Our simulations revealed enhanced transporter activity and more complex metabolic interactions within host-associated microbial communities compared to in vitro conditions (**Fig. 3 F,G; 4 A,B**). Notably, the connectivity of the community metabolic network was markedly higher in host-associated communities, suggesting more intricate interdependencies in the intestinal environment. Cross-feeding is common between members of bacterial communities^72^, such as the gut microbiome^73^, even though its implications for community resilience are still under debate^74^. Notably, pairwise interactions of the OMM^12^ community members were shown to be predominantly negative (only one instance of commensalism was observed in 66 pairs)^26^, despite this, community culture of the OMM^12^ consistently finds most species to coexist^7,18,20^. Our modelling approach suggests that in the community context, exchanges of metabolites between members are common, especially in vivo. Exchanged metabolites can be small molecules of central carbon metabolism, such as pyruvate, malate, or fumarate, amino acids, polysaccharide breakdown products, as well as nucleotides. Of note, the latter have also recently been shown to be important in the interaction between microbiome and the host across the host’s lifespan^75,76^. Pyruvate is a common product of overflow metabolism in various microbes and is excreted or consumed depending on the growth phase^77^.

Extracellular pyruvate can be sensed and imported by specific systems and has wide-ranging effects on nutrient scavenging and stress survival^78,79^. The computational simulation approach allowed us to deeply probe the mechanism leading to this enhanced exchange of metabolites. It revealed that this can be a direct consequence of a change in carbon sources, potentially explaining how bacterial interactions can change from positive to negative between different media as observed by Weiss et al^7^. Pérez Escriva et al. inferred a predicted network of exchanges within the OMM^12^ in vitro, based on metabolite production and consumption^27^. Our modelling approach identified some of the same species as “producer” and “consumer” hubs. *B. caecimuris* I48 was identified by both approaches as a central member of the network, likely making carbon sources available to others, especially in vivo. Species with a broad substrate range and high carbon source adaptability, such as *E. clostridioformis* and *C. innocuum* I46, are also central members of the microbial network and primarily consume many different exchanged metabolites (**Fig. 5B, C; Fig. S18**).

One limitation of our study is the reliance on a small synthetic community with limited complexity. As a result, interactions between very closely related species could not be studied and the environment of the gut, with hundreds of species with overlapping nutrient profiles, is not sufficiently represented. On the other hand, this reduced complexity allowed us to recover proteomic signatures from most members in a depth that allowed predictions about main metabolic functions. Furthermore, the lack of competition between closely related species or species with similar nutrient profiles allowed effects to be read out with more clarity. A second shortcoming is the current limitation of genome-based metabolic models for non-model organisms. Since GEMs rely on homology, they have considerable blind spots in poorly annotated genomes that could only be partially addressed by gap-filling. Also, in our proteomic data, some highly regulated proteins were not functionally annotated.

Notably, large-scale analysis of variable expression of unannotated proteins can help uncover their functions^80^ and represents a future direction for OMM^12^ research. Furthermore, limited knowledge of kinetics and energetic constraints can hinder correct predictions of metabolic models^81^. Still, large-scale applications^45,46,48,56^ have shown that metabolic modelling can correctly recapitulate ecosystem functions and serve as a prediction tool for exchanges of interest^42–46,76^.

In conclusion, our findings highlight the remarkable adaptability of microbial communities across environmental contexts. This underscores the importance of studying microbiomes under conditions that closely approximate their natural habitats and integrating context-specific data into microbiome function studies. The observed shift toward resource acquisition strategies in host-associated communities represents an ecological adaptation to the competitive and complex intestinal environment. This comprehensive understanding of context-dependent community functioning provides valuable insights for future efforts to manipulate and engineer microbial communities for therapeutic or biotechnological applications.

## Methods

### Bacterial strains

Strains used in this study were *Enterococcus faecalis* KB1 (DSM32036), *Bifidobacterium animalis* YL2 (DSM26074), *Acutalibacter muris* KB18 (DSM26090), *Muribaculum intestinale* YL27 (DSM28989), *Flavonifractor plautii YL31* (DSM26117), *Enterocloster clostridioformis* YL32 (DSM26114), *Akkermansia muciniphila* YL44 (DSM26127), *Turicimonas muris* YL45 (DSM26109), *Clostridium innocuum* I46 (DSM26113), *Bacteroides caecimuris* I48 (DSM26085), *Limosilactobacillus reuteri* I49 (DSM32035), and *Blautia coccoides* YL58 (DSM26115). Further information and metabolic characteristics can be found in **Table S10**.

### Cultivation and Media

Single bacteria as well as OMM^12^ communities were cultivated in AF and APF as described in ^7,26^, the addition of 0.5g/l extra glucose was omitted from AF medium. Cultures were grown in 24 well plates for 24h at 37°C without shaking in an anoxic atmosphere (7% H_2_, 10% CO_2_, 83% N_2_), with the exception of *A. muris* KB18 and *T. muris* YL45 monocultures, which were grown for 48h before harvest. OMM^12^ communities were assembled fresh from monocultures that were transferred once after inoculation from frozen stocks. Monocultures were diluted to OD 0.1 and then mixed in equal amounts. The resulting community was transferred 5 times into fresh medium before harvest for qPCR and proteomic sample generation.

### DNA extraction and quantitative PCR

For DNA extraction, 1 ml of culture was centrifuged at 14 000 rpm for 2 minutes at 4°C, the supernatant was removed and the pellet stored at -20°C until extraction. Phenol/chloroform extraction followed by DNA purification using a NucleoSpin gDNA clean-up kit (Macherey–Nagel) was performed as described in^26^. Quantitative qPCR on the 16S rRNA gene with strain-specific primers was performed and analysed as in ^25^ with 5 ng of genomic DNA as template on a Roche Diagnostics “LightCycler” with the corresponding software “LightCycler 96” (Version 1.1.0.1320).

### Proteomic sample generation

For proteomic data analysis, 10-20ml (50 ml for *T. muris* YL45 and *A. muris* KB18 monocultures) of an in vitro culture in late exponential/early stationary phase (**Table S1**) were centrifuged (2 min, 12 000 *g*, 4°C), washed twice with the same volume of ice-cold, sterile phosphate-buffered saline, snap frozen in liquid N_2_ and stored at -80°C until sample extraction. Stabilised in vitro OMM^12^ communities were harvested 24 hours after inoculation as described above. For the mouse samples, C57Bl/6 mice stably colonised with the OMM^12^ community were housed under sterile conditions in flexible film isolators (North Kent Plastic Cages) and supplied with autoclaved ddH2O and Mouse-Breeding complete feed for mice (Ssniff) ad libitum as described in^7^. For sample generation, 22 week old male mice were sacrificed by cervical dislocation and cecum content was harvested. Cecum content was resuspended in 20 ml of ice-cold, sterile phosphate-buffered saline and filtered through a 40 µm cell strainer (Greiner Bio-One EASYstrainer) to remove debris. 6ml of the flowthrough were centrifuged (2 min, 12 000 *g*, 4°C), snap frozen in liquid N_2_ and stored at -80°C until sample extraction.

Bacteria were lysed with 100% trifluoroacetic acid (TFA) according to the SPEED protocol^82^ with slight adaptations^83^. Briefly, 100µl of trifluoroacetic acid were added to each bacterial pellet and incubated at 55°C for 5 minutes. Samples were neutralised by the addition of 900µl 2M TRIS and stored at -20°C until analysis. Protein concentrations were determined by Bradford assay according to manufacturer’s instructions against a BSA standard and a negative control of 10% trifluoroacetic acid in 2M TRIS.

Next, 50 µg total protein amount per sample was reduced and alkylated (9 mM tris(2-carboxyethyl)phosphine (TCEP) and 40 mM chloroacetamide (CAA) for 5 min at 95 °C) in one step. Samples were diluted with deionized water to a final concentration of 1 M Tris and 5% TFA. Trypsin (Roche), was added in a trypsin:protein ratio of 1:50. Samples were incubated overnight at 37 °C and 400 rpm. Next day, all samples were acidified to a final concentration of 3% formic acid (FA) and desalted. Briefly, self-packed desalting tips were prepared in-house, consisting of five Empore C18 (3M). Tips were primed with 100% acetonotrile (ACN), then 40% ACN/0.1% FA, and finally equilibrated with 0.1% FA. Peptides were loaded, washed with 0.1% FA, and eluted twice with 40% ACN/ 0.1% FA. Samples were lyophilized and dried peptides were stored at –80 °C.

### Proteomic data acquisition

Peptides were analyzed on a Vanquish^TM^ Neo UHPLC (microflow configuration; Thermo Fisher Scientific, MA, USA) coupled to an Orbitrap Exploris^TM^ 480 mass spectrometer (Thermo Fisher Scientific, MA, USA). Around 25 µg of peptides were applied onto a commercially available Acclaim PepMap 100 C18 column (2 μm particle size, 1 mm ID × 150 mm, 100 Å pore size; Thermo Fisher Scientific, MA, USA) and separated on a stepped gradient from 3% to 31% solvent B (0.1% FA, 3% DMSO in ACN) in solvent A (0.1% FA, 3% DMSO in HPLC grade water) over 60 min. A flow rate of 50 μl/min was applied. The mass spectrometer was operated in DDA and positive ionization mode. MS1 full scans (360 – 1300 m/z) were acquired with a resolution of 60,000, a normalized automatic gain control target value of 100%, and a custom maximum injection time of 50 ms. Peptide precursor selection for fragmentation was carried out using a cycle time of 1.2 seconds. Only precursors with charge states from two to six were selected, and dynamic exclusion of 30 s was enabled. Peptide fragmentation was performed using higher energy collision-induced dissociation and a normalized collision energy of 28%. The precursor isolation window width of the quadrupole was set to 1.1 m/z. MS2 spectra were acquired with a resolution of 15,000, a fixed first mass of 100 m/z, a normalized automatic gain control target value of 100%, and a custom maximum injection time of 40 ms.

### Proteomic data analysis

For proteomic analysis, MaxQuant v1.6.17.0 with its built-in search engine Andromeda^84,85^ was used. MS2 spectra were searched against combined fasta files of all 12 members of the OMM^12^ community, including *E. faecalis* KB1, *F. plautii* YL31, *C. innocuum* I46, *B. animalis* YL2, *E. clostridioformis* YL32, *B. caecimuris* I48, *A. muris* KB18, *A. muciniphila* YL44, *L. reuteri* I49, *M. intestinale* YL27, *T. muris* YL45 and *B. coccoides* YL58. Their respective genomes were obtained from GenBank and RefSeq (accession numbers see **Table S10**)^86^. GenBank genomes were used as a template and unique proteins from RefSeq added based on their coordinates. KEGG annotations and KOs were added using KofamKOALA^87^ (https://www.genome.jp/tools/kofamkoala/). Proteomes were obtained from these genomes and supplemented with common contaminants (built-in option in MaxQuant). Trypsin/P was specified as the proteolytic enzyme. Precursor tolerance was set to 4.5 ppm, and fragment ion tolerance to 20 ppm. The minimal peptide length was defined as seven amino acids. The “match-between-run” function was disabled. Carbamidomethylated cysteine was set as a fixed modification, and methionine oxidation and N-terminal protein acetylation were defined as variable modifications. Results were adjusted to a 1% false discovery rate on peptide spectrum match (PSM) level and protein level, employing a target-decoy approach using reversed protein sequences. Label-Free Quantification (LFQ)^88^ intensities were used for protein quantification with at least 2 peptides per protein identified. Only proteins identified in at least two out of three biological replicates were considered and no values were imputed. Data was analysed and volcano plots and bar charts were generated in GraphPad Prism (10.3.1) and R (4.4.2) using the tidyverse set of packages (version 2.0.0).

Hierarchical clustering was performed on unnormalised protein intensities in R (4.4.2) using the dist function to calculate Euclidean distances and hclust with complete agglomeration and plotted using ggdendro (version 0.2.0) and ggplot2 (version 3.5.1).

For the analysis of monoculture vs. community culture in AF, protein signals were separated by species and ranked within each species-specific proteome. Rank differences were analysed with unpaired t-tests and multiple comparisons were corrected for using a false discovery rate approach with a q-value cutoff < 0.05 using the two-stage step-up method by Benjamini, Krieger, and Yekutieli^51^.

For the comparison of OMM^12^ communities grown in different media, protein signals were separated by species and divided by the total sum of MS intensities for the species in the given sample. Differences were analysed with unpaired t-tests and multiple comparisons were corrected for using a false discovery rate approach with a q-value cutoff < 0.05 using the two-stage step-up method by Benjamini, Krieger, and Yekutieli.

### Functional analysis

Proteins with a rank change > 100 in either direction in the comparison between monoculture and community culture were considered for enrichment. These proteins were subjected to enrichment analysis using the enrichKEGG function in the R package clusterProfiler (version 4.6.2)^89^. Enrichment analysis between community cultivation conditions was performed with the same function on all proteins with a significant difference (FDR < 0.05) between conditions and a fold change of 2 in either direction, as well as all proteins exclusive to one condition with a normalized signal of more than 0.0001 (i.e. that contribute more than 0.01% of total cell protein).

For the analysis of proteome allocation, proteins were sorted into COG categories using eggNOG-mapper v2^90^ and compared between growth conditions. Comparison of monoculture versus community culture was analysed with unpaired t-tests and multiple comparisons were corrected for using a false discovery rate approach with a q-value cutoff < 0.05 using the two-stage step-up method by Benjamini, Krieger, and Yekutieli. Signals from different communities (AF, APF, mouse) were compared with a two-way ANOVA followed by pairwise comparisons corrected by Tukey’s test (confidence interval 95%).

Signal peptides were predicted by SignalP v4.1^91^ using the dbCAN v4^92^.

### Metabolic model generation

We used available genomes^86^ of each member of the OMM^12^ community to reconstruct metabolic network models using gapseq (1.4.0 88d4b682)^53^ with a bitscore of 200 for reaction predictions. For the in silico growth medium, we build on our previously published growth medium^26^, which was slightly adapted to match the AF and APF medium used in experiments. After the automated reconstruction process, we addressed inconsistency in the resulting models. All changes were integrated into gapseq, ensuring that shortcomings are also addressed for future model building. In particular, we updated protein complex handling in gapseq (a0b1380), available sequences for propanediol hydratase (6be0b6b), inulin degradation (72ff3e0), and lysine (098aee6) and arginine degradation (4b6b6f3) (the codes correspond to SHA hashes of commits, see https://github.com/jotech/gapseq/commits for details). The models are available as SBML files (https://doi.org/10.5281/zenodo.17358311). The logistic PCA in Figure 4a was done by using the R package logisticPCA described in^93^.

### Generation of context-specific models

We refined the metabolic models created with gapseq by considering those enzymes supported by proteomic data. For this, we made a list of core reactions for each organism and condition by matching experimentally measured proteins to genes and genes to model reactions. We assumed each protein with a non-zero measurement to be present. Also, we included reactions with no associated genes (e.g., biomass reaction and exchange reactions) in the core reaction lists. The core list was then used as input for fastcore^94^ to obtain a refined metabolic network model consisting of a minimized amount of reactions supported by experimental evidence and overall network structure. We used the fastcore implemented in Python by the troppo package (https://github.com/BioSystemsUM/troppo) as described previously^95^. The context-specific models are available as SBML files (https://doi.org/10.5281/zenodo.17358311).

### Validation of metabolic model predictions

We used experimental data from Weiss et al.^26^ and Pérez Escriva et al.^27^ to verify our metabolic models for monocultures. Known uptake and production capabilities of amino acids (18), carbohydrates (2), fatty acids (3), and organic acids (2) for all 12 members of the OMM^12^ community were compared to model predictions. For this pFBA was performed and exchange rates compared to experimental values. We compared three states, production (indicate by +1), uptake (−1), and no change (0) (see **Table S6**). For metabolites which could not be distinguished experimentally (L-lactate/D-Lactate and leucine/isoleucine), we took the mean of simulated fluxes for comparison. We only considered amino acids uptake vor validation because amino acids could also emerge due to degradation of larger proteins. In two cases for YL58, the uptake of asparagine and histidine was unsure, we therefore removed the two cases. As performance metrics, we used accuracy, (TP+TN)/(TP+TN+FP+FN), and kappa as implemented in the caret R package^96^ (see script *validation_mono.r* for more details).

Short-chain fatty acid and amino acid profiles of in vitro (AF and APF) and in vivo communities were compared against data for OMM^12^ communities from Weiss et al^7^. Simulated values for D- and L-lactate were added together to compare to the experimentally determined lactate signal and simulated leucine and isoleucine values were added to compare to the experimentally determined leucine/isoleucine signal. Simulated values were extracted from the *medlist* component of the Eval object for each simulation and values at timepoint 1 were subtracted from values at timepoint 4. Experimental values were calculated with medium blanks subtracted from in vitro community measurements. In vivo concentrations were used as is since no blank was available. Plotting was performed in Prism (10.3.1).

### Community modelling and network analysis

Community growth using context-specific models was modelled in BacArena V1.8.1^56^. Relative qPCR data matching the metaproteomic data from OMM^12^ communities grown in AF and APF, as described above and extracted from the mouse cecum, were used to determine the starting distribution of OMM^12^ members. APF was used as the medium to model the mouse gut condition after cross-referencing with published metabolites from the germ-free mouse gut^97^.

Metabolic networks were extracted using the FindFeeding3 function in BacArena and plotted using the igraph package V2.1.2^98^. The degree of connection for each node was extracted using the degree function in igraph and plotted using Prism (10.3.1)

### Figures

Figures 1A and 2A were generated using BioRender (https://biorender.com). All Figures were composed using Affinity Designer 2 (Version 2.6.3).

## Code and data availability

The mass spectrometric raw files as well as the MaxQuant output files have been deposited to the ProteomeXchange Consortium via the PRIDE partner repository. They can be accessed using the identifier PXD069036 (https://proteomecentral.proteomexchange.org/cgi/GetDataset?ID=PXD069036).

Code used in this study is available at https://github.com/annaburrichter/OMM12-modelling and the metabolic models can be found at https://doi.org/10.5281/zenodo.17358311.

## Acknowledgements

The authors thank Diana Ring for excellent technical assistance in cultivation, Monica S. Matchado for help with data analysis and Hermine Kienberger, Verena Breitner and Franziska Hackbarth for excellent technical assistance and maintenance of mass spectrometers. B.S. and A.G.B. received funding by the European Research Council (ERC) under the European Union’s Horizon 2020 research and innovation program (grant agreement 865615) and B.S. received additional funding by the German Research Foundation (DFG, German Research Foundation, SFB 1371, project number 395357507). J.Z. acknowledges support from the Deutsche Forschungsgemeinschaft (DFG, German Research Foundation) under Germanýs Excellence Strategy – EXC 2051 – Project-ID 390713860. The Exploris 480 mass spectrometer was funded in part by the German Research Foundation (INST 95/1435-1 FUGG).

## Author contributions

A.G.B. and J.Z. conceived and wrote the manuscript. A.G.B. designed and performed the experiments and analyzed data. J.Z. created and analyzed metabolic and context-specific models. A.G.B. and J.Z. performed community simulations and interpreted results. C.L. and C.M. acquired and analyzed proteomic data. B.S. and C.K. supervised the project. All coauthors provided feedback and reviewed the manuscript.

## Supplementary Data

Figure S1: relative abundances of OMM^12^ members in the community for the monoculture - community comparison.

Figure S2: Proteomic data of monocultures compared to strain-resolved community culture.

Figure S3: KEGG enrichment over all strains of all proteins between monoculture and strain-resolved community culture.

Figure S4: KEGG enrichment by strain of all proteins between monoculture and strain-resolved community culture.

Figure S5: Expression levels of potential extracellular proteins.

Figure S6: abundances of OMM^12^ members of community for in vitro - in vivo comparison as determined by qPCR and protein intensities.

Figure S7: Expression data of specific pathway enzymes in OMM^12^ communities grown in glucose-rich (AF) and polysaccharide-rich (APF) medium as well as in the mouse cecum.

Figure S8: Volcano plots of proteomic data AF vs. APF and AF vs. mouse.

Figure S9: KEGG enrichment over all species between cultivation conditions (in vitro-AF medium, in vitro-APF medium, mouse cecum).

Figure S10: KEGG enrichment per species of all proteins between cultivation conditions (in vitro-AF medium, in vitro-APF medium, mouse cecum).

Figure S11: proteome allocation per COG category for single species in monoculture vs. community culture.

Figure S12: proteome allocation by COG category groups by species in in vitro and host-associated communities.

Figure S13: Fractions of community proteome (all species) by COG category under each community cultivation condition.

Figure S14: context-specific model coverage.

Figure S15: Comparison of metabolic network sizes of context-specific models. Figure S16: Similarity of condition-specific metabolic models between conditions.

Figure S17: Predicted growth curves of the OMM^12^ community with AF (A), APF (B) medium and in mouse (C).

Figure S18: Connectivity of species within the community network broken down into ingoing and outgoing edges.

Figure S19: Comparison of simulated short-chain fatty acid and amino acid production and consumption data with experimental data.

Table S1: cultivation time for proteomic samples

Table S2: Data points marked in volcano plots (Figures 1D-H and S2)

Table S3: full datasets for enrichment monoculture – community

Table S4: full datasets for enrichment in vitro - in vivo

Table S5: Full gene annotations of community expression data (Figures 2F and S6)

Table S6: reference data for model validation (Figures 2D and S19)

Table S7: predicted exchange fluxes in the metabolic network models (Figure 4B)

Table S8: reactions involving pyruvate in the I46 mouse CSM (Figure 4D)

Table S9: expression of PTS system components in C. innocuum I46 and of the trehalose PTS system in E. faecalis KB1

Table S10: Strain characteristics and identifiers for OMM^12^

