## Supplementary Figures and Tables for "Context-Dependent Metabolic Adaptation in Microbial Communities: From Monocultures to Complex Ecological Interactions"

Table S7: predicted pyruvate exchange fluxes in the mouse metabolic network model (Figure 4B)

Table S8: reactions involving pyruvate in the I46 mouse CSM (Figure 4D)

Table S9: expression of PTS system components in *C. innocuum* I46 and of the trehalose PTS system in *E. faecalis* KB1

Table S10: Strain characteristics and identifiers for OMM[12]

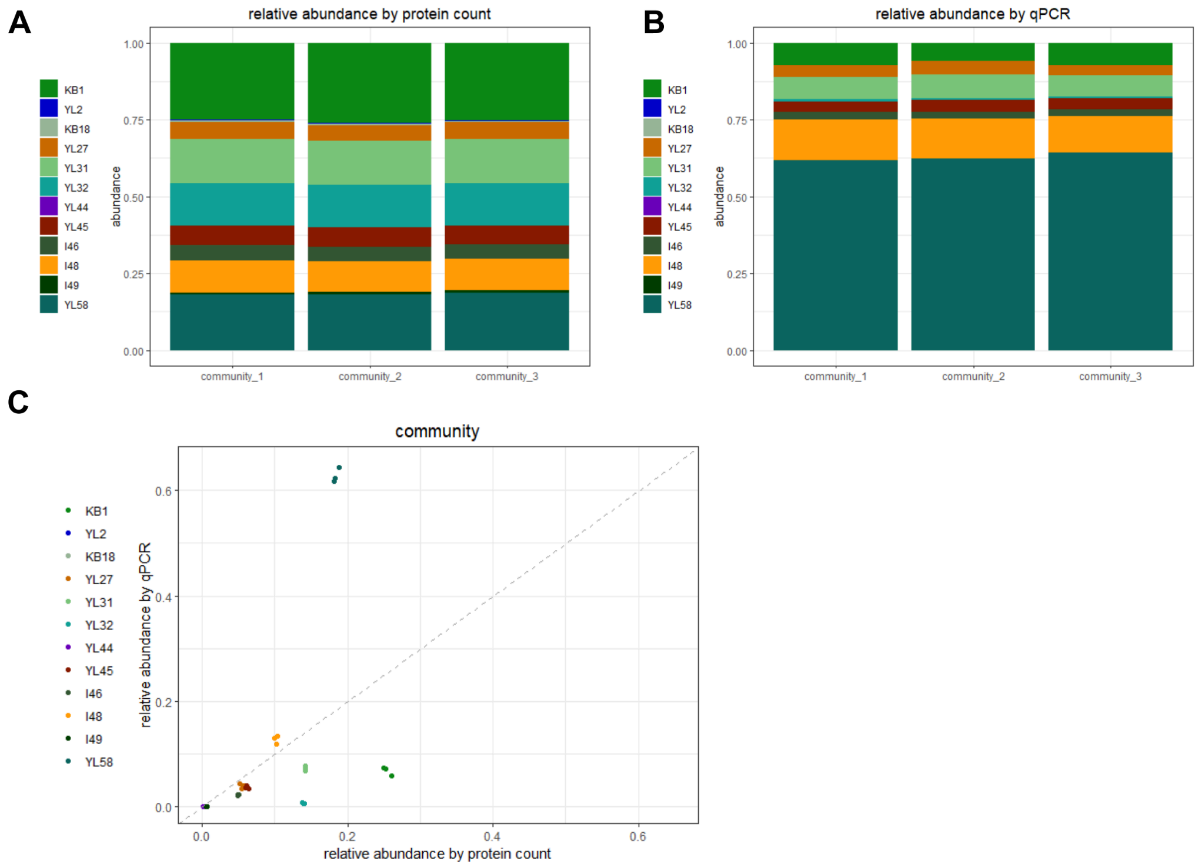


**Figure S1: relative abundances of OMM$^{12}$ members of community for monoculture - community comparison as determined by protein count (A) and qPCR (B) and the correlation between both measurements.**

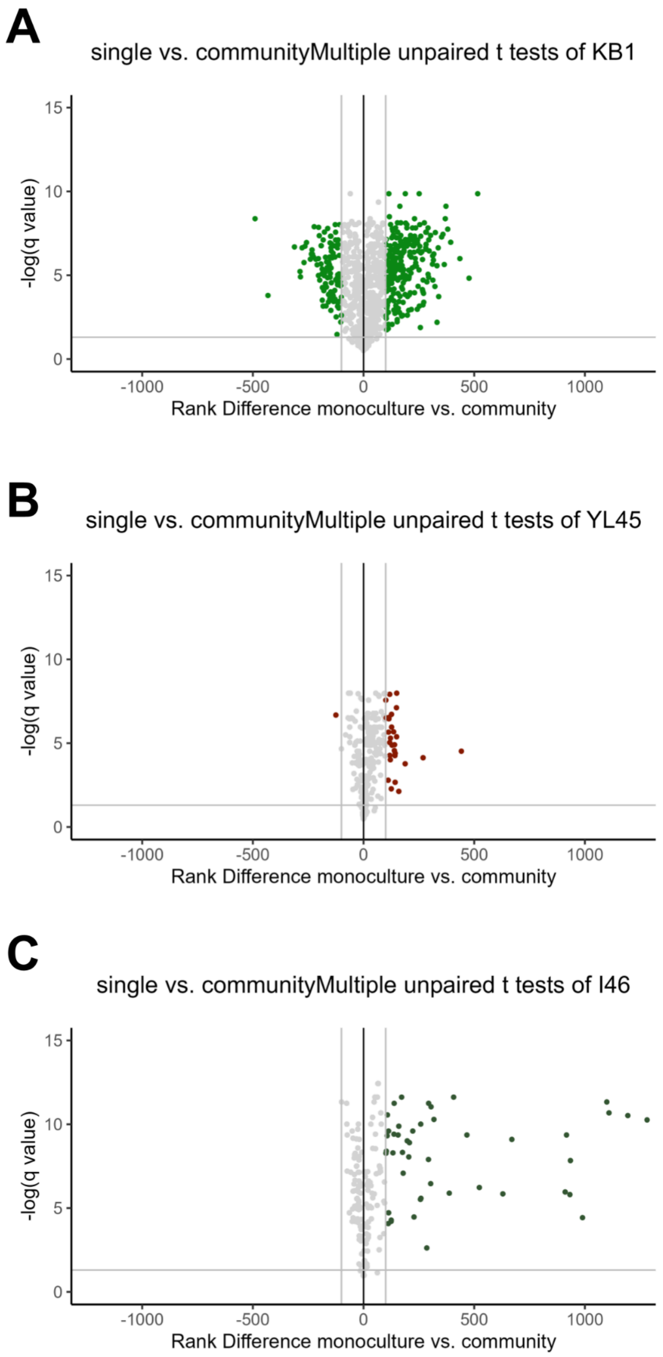


**Figure S2: Proteomic data of monocultures compared to strain-resolved community culture for *E. faecalis* KB1 (A), *T. muris* YL45 (B), and *C. innocuum* I46 (C). Volcano plots of protein rank in monoculture versus strain-resolved community culture show proteins overexpressed in the community context on the right hand side of each plot. x axis: rank difference monoculture - community culture; y axis: q value of unpaired multiple t-tests with a false discovery rate correction for multiple comparisons (FDR cutoff 1% with the two-stage step-up method of Benjamini, Krieger and Yekutieli); grey lines mark a rank change of 100 and a q value of 0.05.**

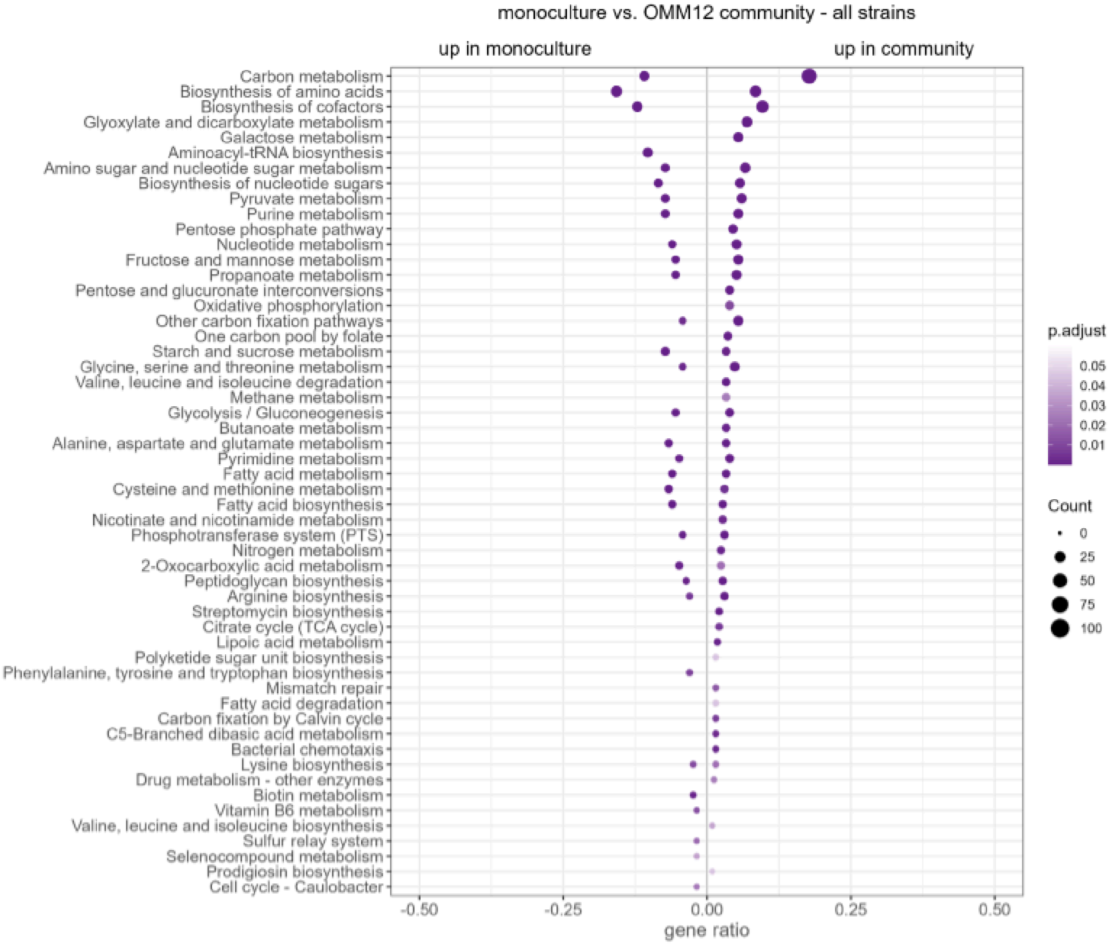


**Figure S3: KEGG enrichment over all strains of all proteins with a significant rank change > 100 in their respective species-specific proteome between monoculture and strain-resolved community culture in AF medium.**

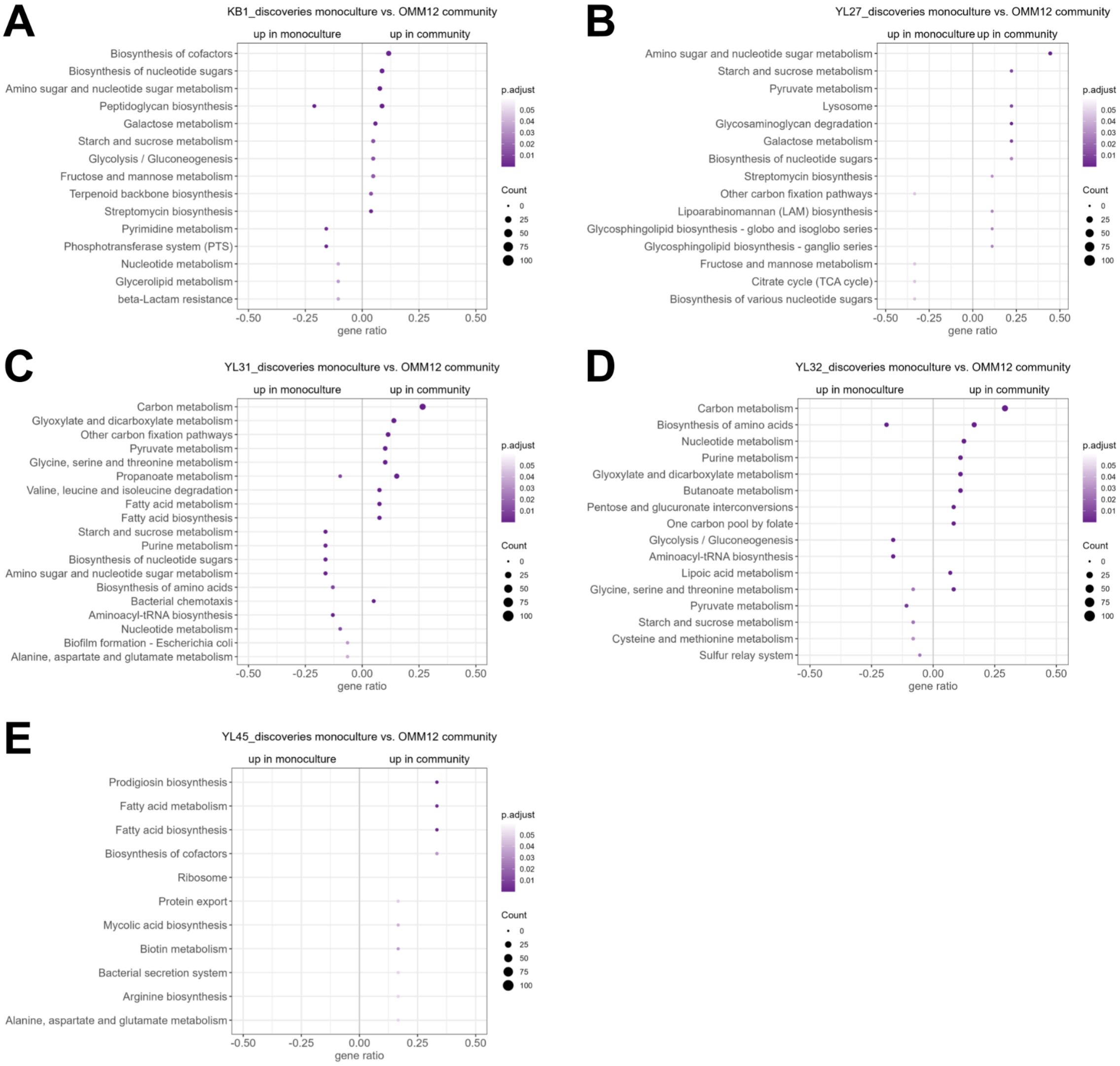


**Figure S4: KEGG enrichment by strain of all proteins with a significant rank change > 100 in their respective species-specific proteome between monoculture and strain-resolved community culture in AF mediumfor *E. faecalis* KB1, *M. intestinale* YL27, *F. plautii* YL31, *E. clostridioformis* YL32, and *T. muris* YL45.**

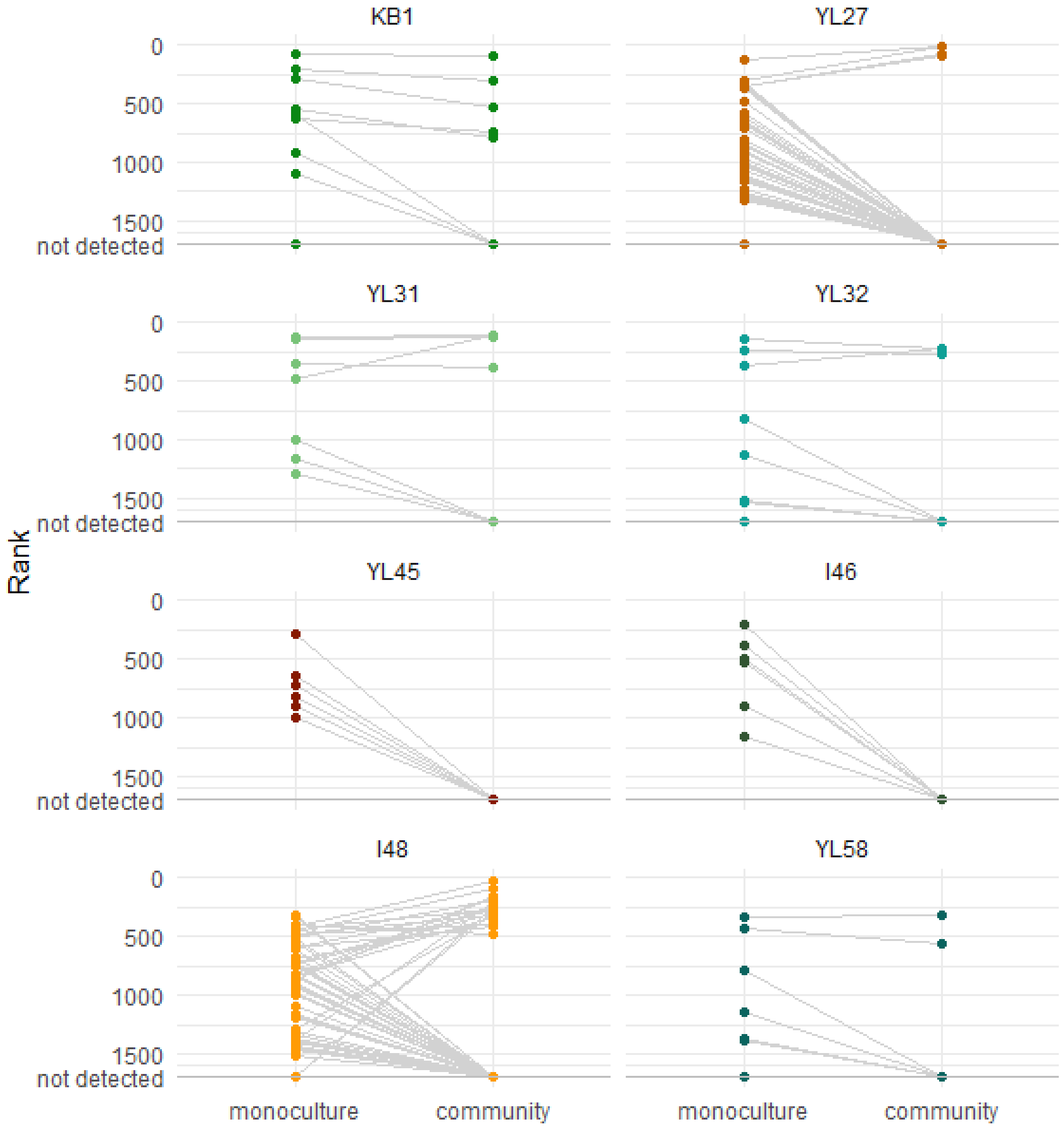


**Figure S5: Expression levels of potential extracellular proteins (containing signal peptides for export) in monoculture versus community culture. Signal peptides were predicted by SignalP v4.1 using the dbCAN v4.**

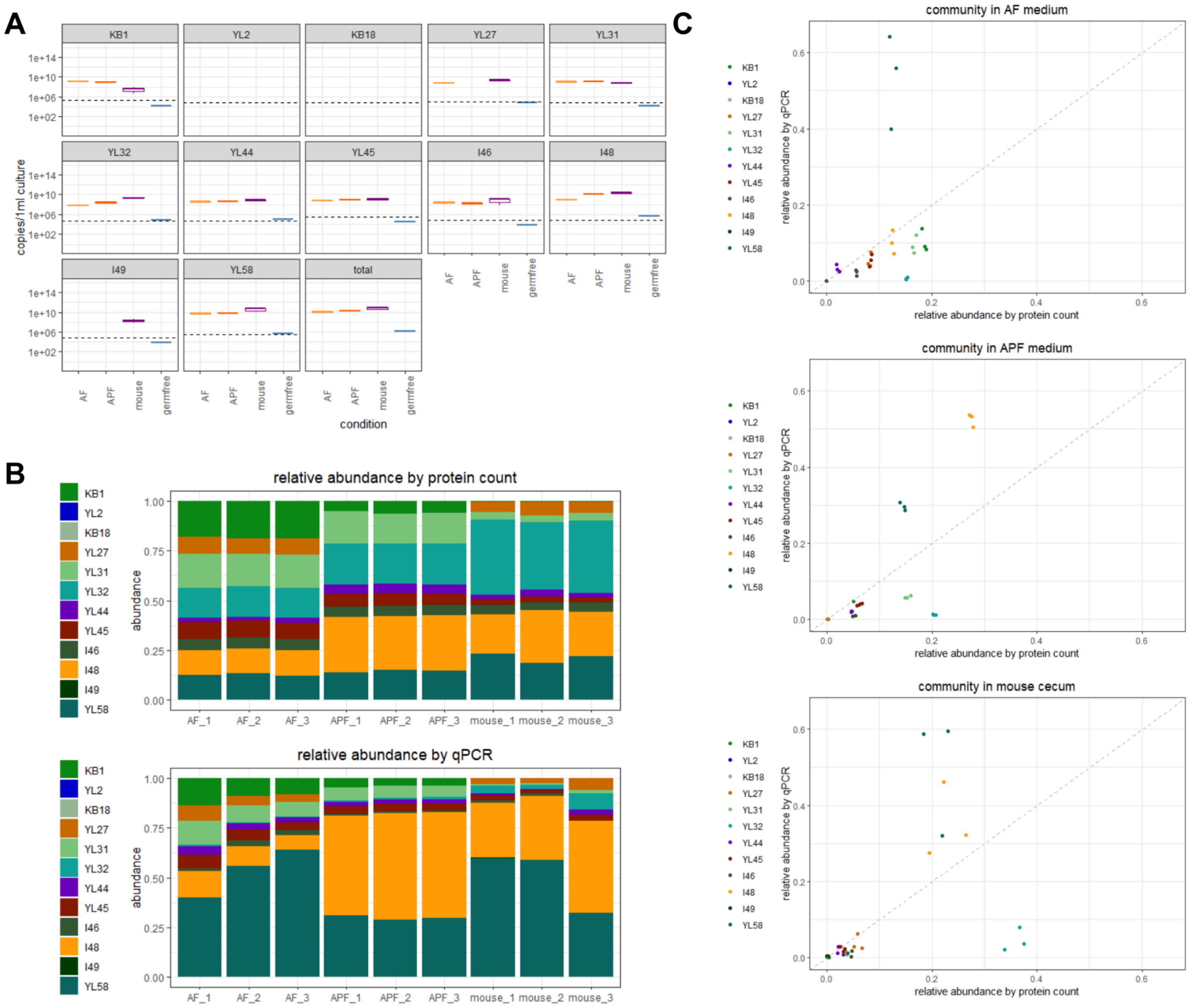


**Figure S6: abundances of OMM$^{12}$ members of community for in vitro - in vivo comparison as determined by qPCR and protein intensities. A: absolute values ( copies of the 16S rRNA gene per ml of liquid culture or mg cecal content, including one sample from a germ-free animal as a control), dotted lines denote the limit of detection. B: relative abundances as determined by protein count (top) and qPCR (calculated from the values in A) (bottom). C: correlation for relative abundances determined by protein count and qPCR for the AF (top), APF (center), mouse (bottom).**

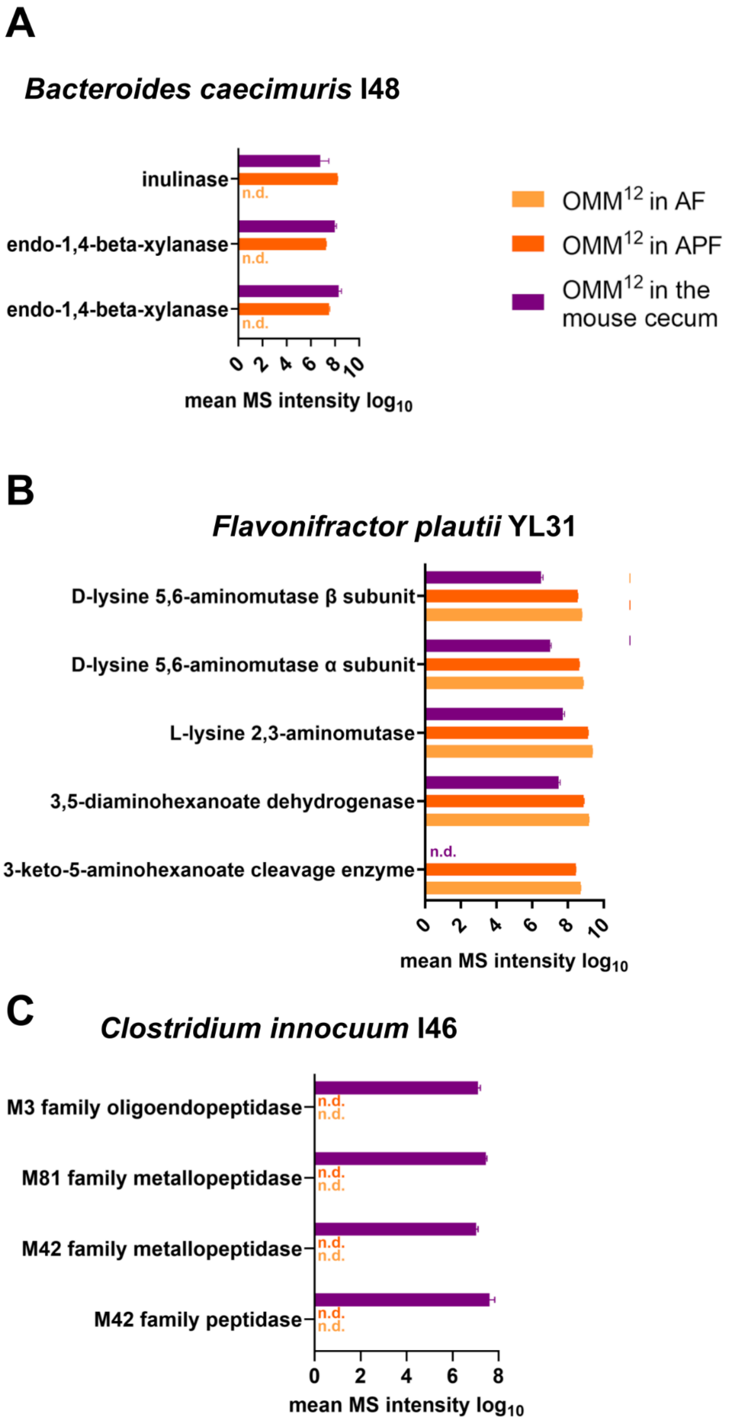


**Figure S7: Expression data of specific pathway enzymes in OMM[12] communities grown in glucose-rich (AF) and polysaccharide-rich (APF) medium as well as in the mouse cecum. A: Inulinase and xylanases of *B. caecimuris* I48. B: Lysine degradation pathway of *F. plautii* YL31. D: Peptidases of *C. innocuum* I46. Full protein annotations can be found in Table S5. n.d.: not detected**

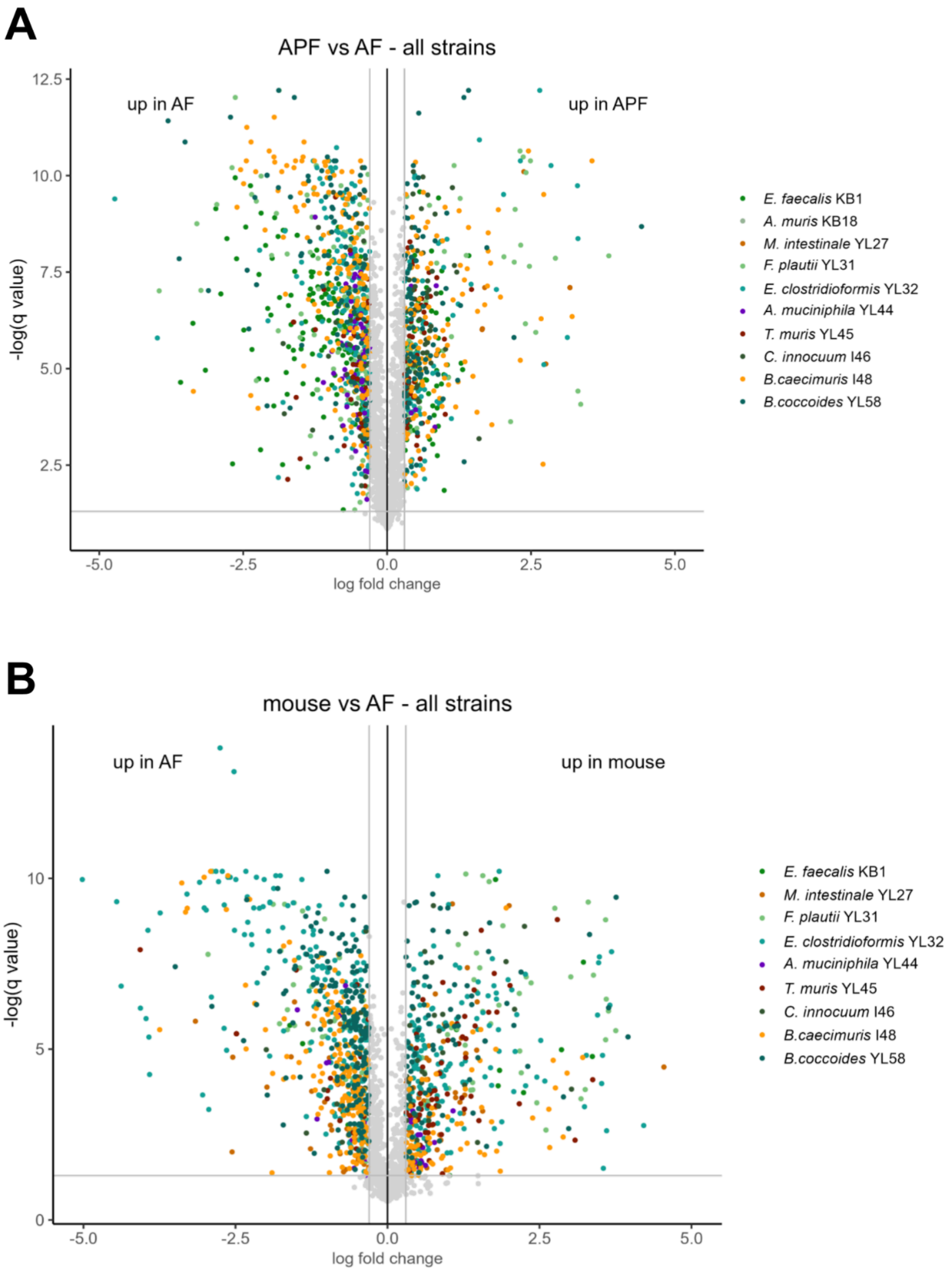


**Figure S8: Volcano plots of proteomic data AF vs. APF and AF vs. mouse. x axis: log fold change of species-normalised protein intensities; y axis: q value of unpaired multiple t-tests with a false discovery rate correction for multiple comparisons (FDR cutoff 5% ; grey lines mark a fold change of 2 and a q value of 0.05.**

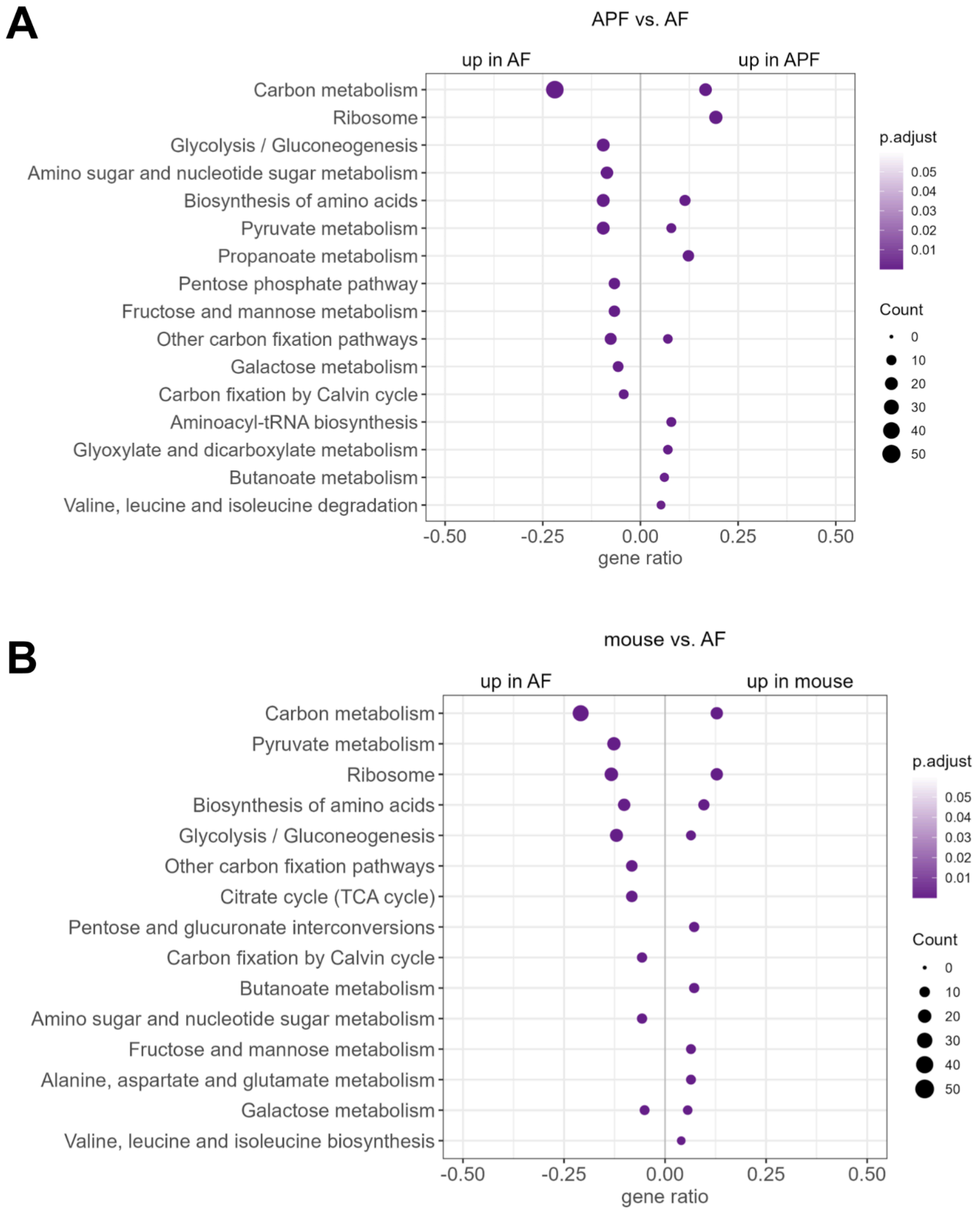


**Figure S9: KEGG enrichment over all species of all proteins with a significant fold change > 2 between cultivation conditions (in vitro-AF medium, in vitro-APF medium, mouse cecum). MS intensity of protein signals was normalised via the total MS intensity of the respective species (i.e., sum of all protein MS intensities assigned to that species). A: AF vs. APF. B: mouse vs. AF.**

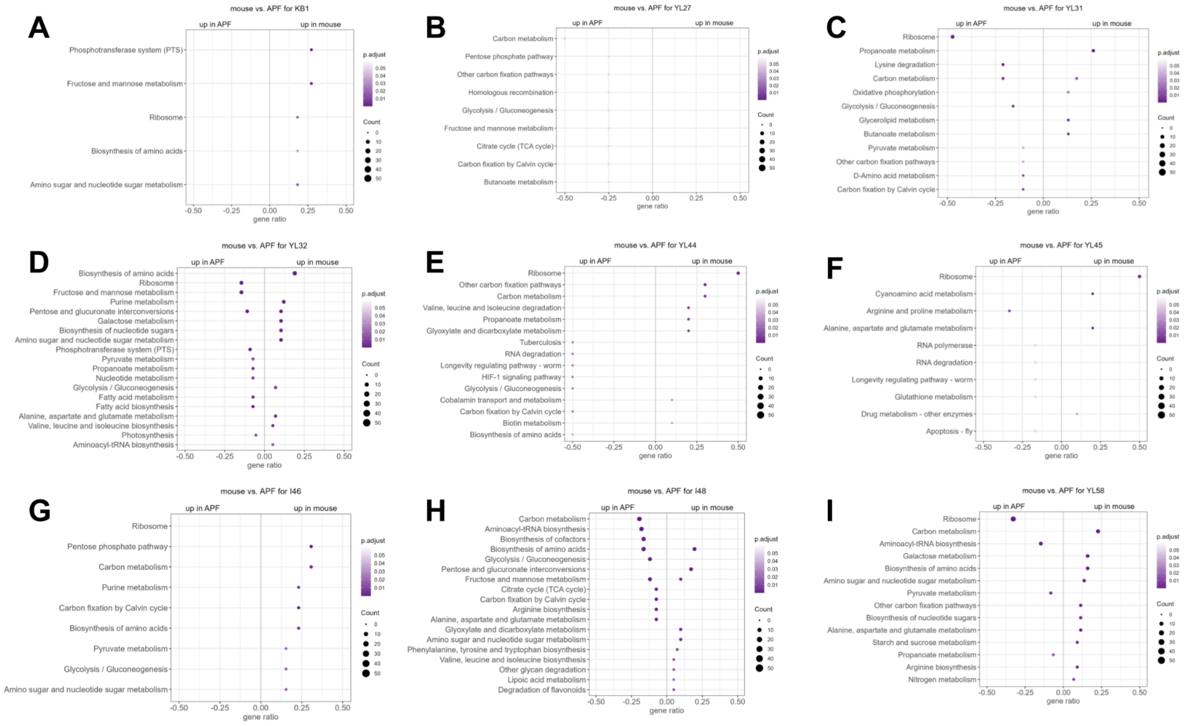


**Figure S10: KEGG enrichment per species of all proteins with a significant fold change > 2 between in vitro-APF medium and mouse cecum. MS intensity of protein signals was normalised via the total MS intensity of the respective species (i.e., sum of all protein MS intensities assigned to that species). A: *Enterococcus faecalis* KB1. B: *Muribaculum intestinale* YL27. C: *Flavonifractor plautii* YL31. D: *Enterocloster clostridioformis* YL32. E: *Akkermansia muciniphila* YL44. F: *Turicimonas muris* YL45. G: *Clostridium innocuum* I46. H: *Bacteroides caecimuris* I48. I: *Blautia coccoides* YL58. No data for *Acutalibacter muris* KB18, *Bifidobacterium animalis* YL2 and *Limosilactobacillus reuteri* I49.**

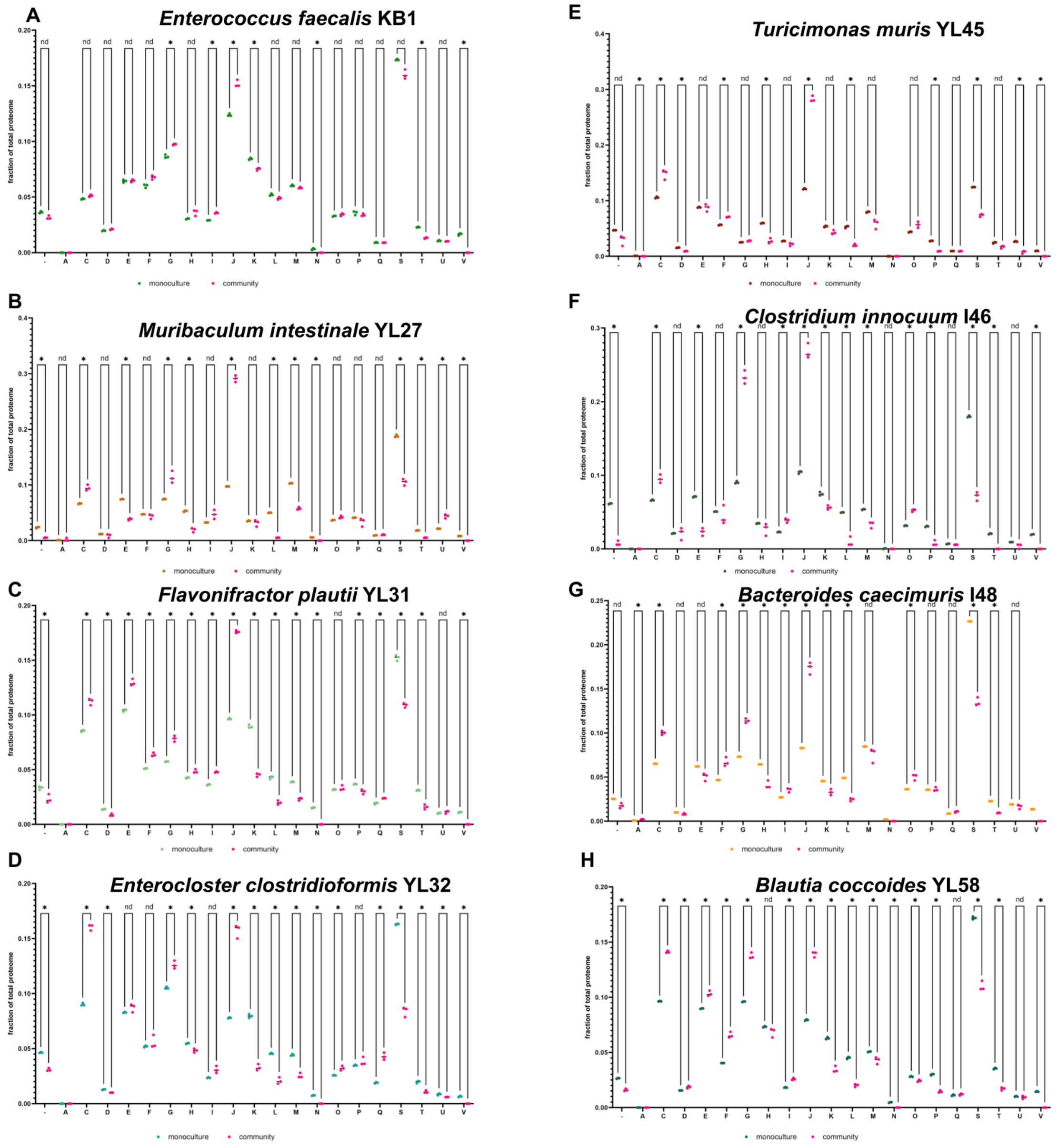


**Figure S11: proteome allocation per COG category for single species in monoculture vs. community culture (statistics: multiple unpaired t-tests with a false discovery rate correction for multiple comparisons (FDR cutoff 0.5% with the two-stage step-up method of Benjamini, Krieger and Yekutieli).**

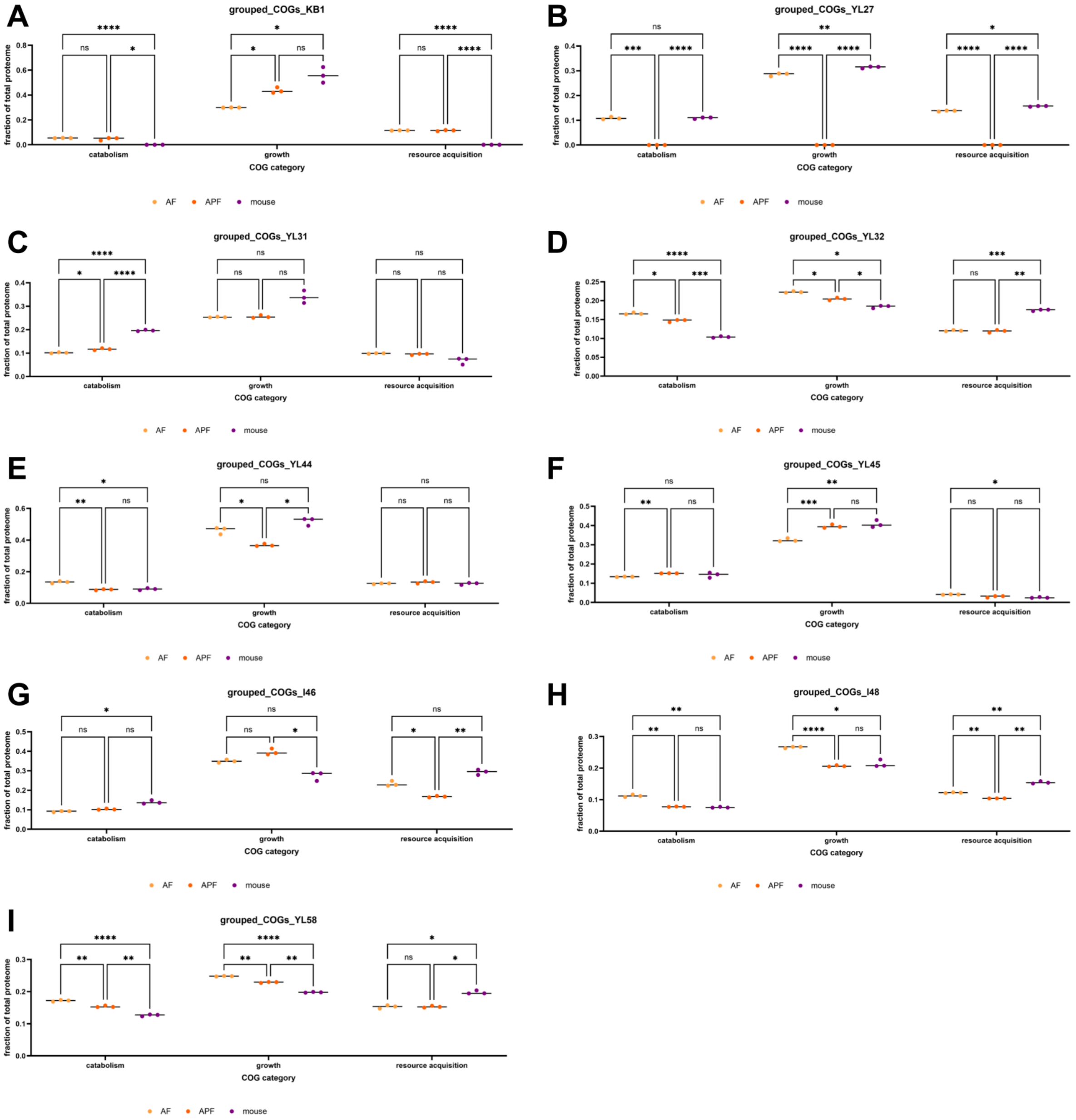


**Figure S12: proteome allocation by COG category groups by species in in vitro and host-associated communities (statistics: two-way ANOVA followed by Tukey's test with confidence interval 95%).**

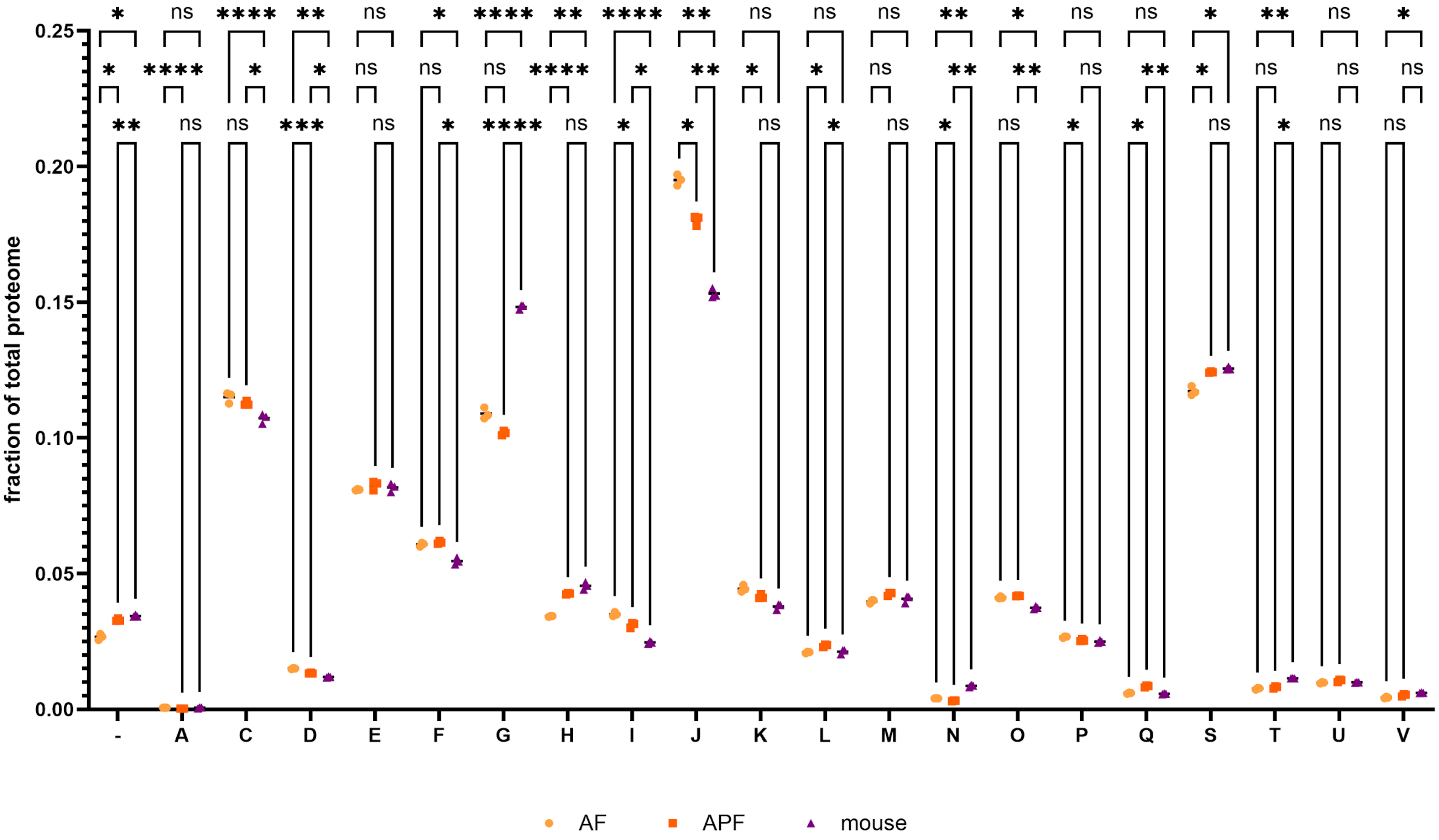


**Figure S13: Fractions of community proteome (all species) by COG category under each community cultivation condition. Statistical analysis: Two-way ANOVA followed by Tukey's multiple comparisons test.**

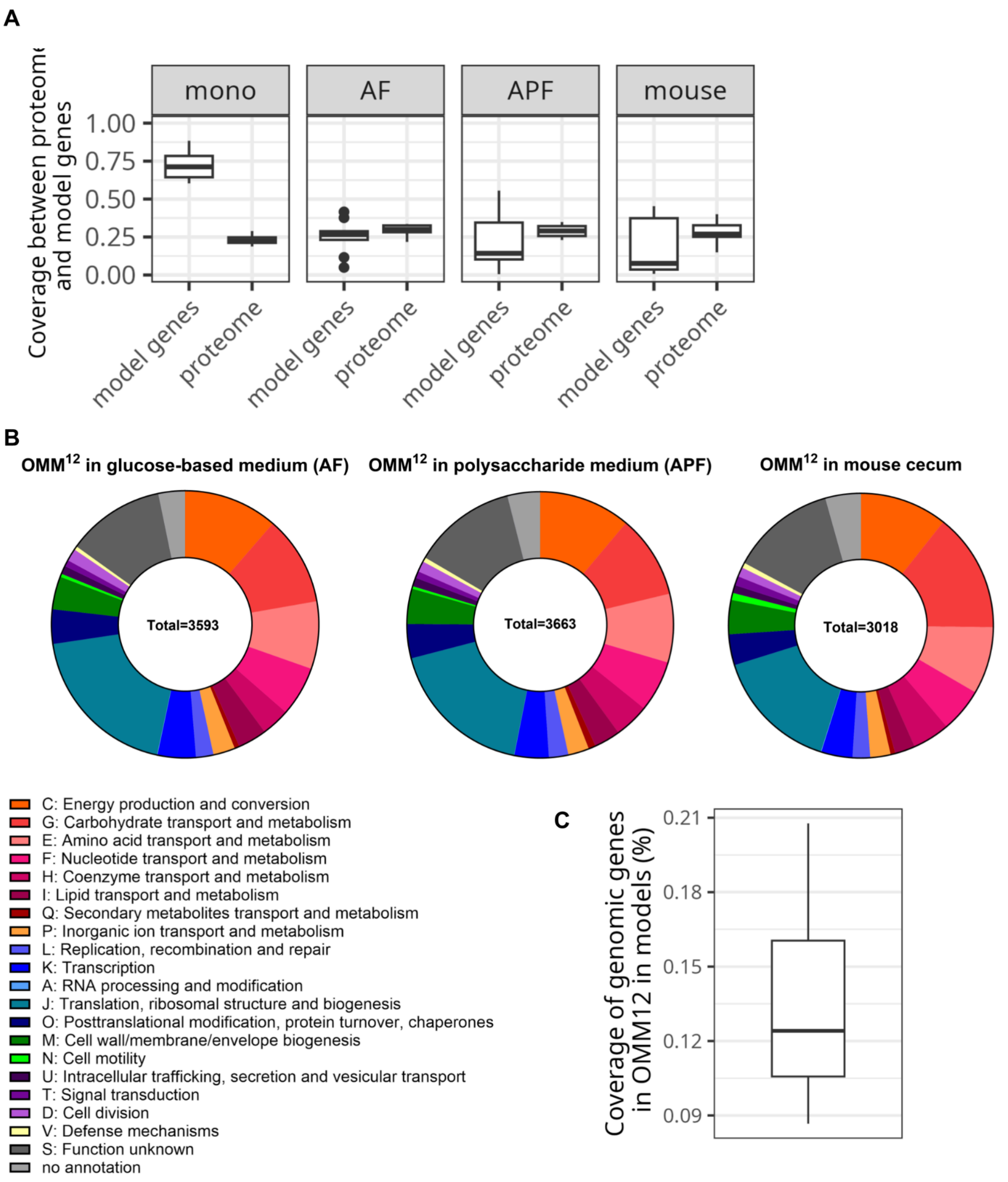


**Figure S14: context-specific model coverage. A: proportion of model reactions that are represented by a measured protein ("model genes") and proportion of measured proteins that are represented by a model reaction ("proteome"). B: COG allocation of measured proteins (red shades represent metabolic categories that can theoretically be represented by a GEM reaction). C: Percentage of genes in the genomes of OMM$^{12}$ members that are linked to reactions in the corresponding metabolic models.**

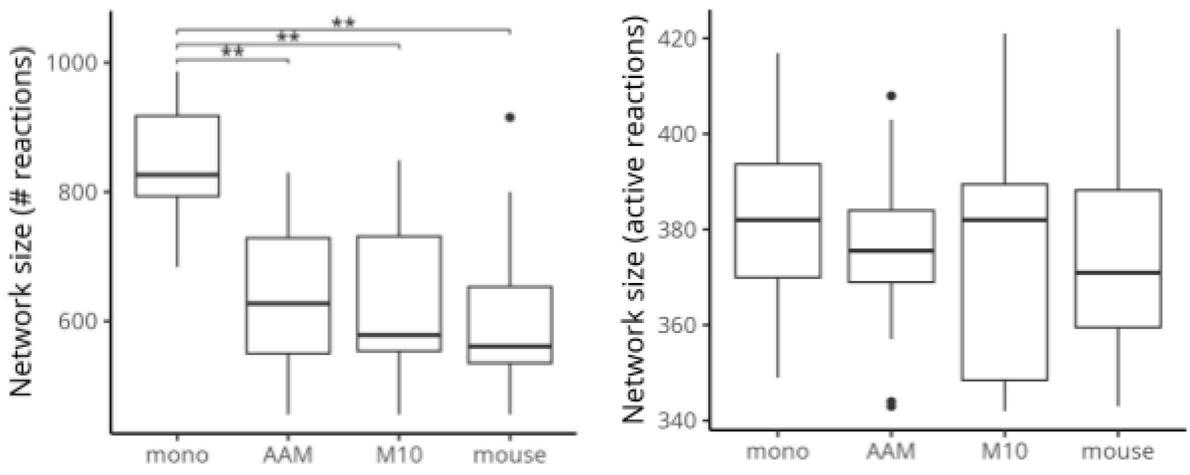


**Figure S15: Comparison of metabolic network sizes of context-specific models. The left subfigure shows the overall reaction count per condition. The right subfigure depicts the number of active reactions (i.e., non-zero fluxes determined by minimizing the total flux optimization).**

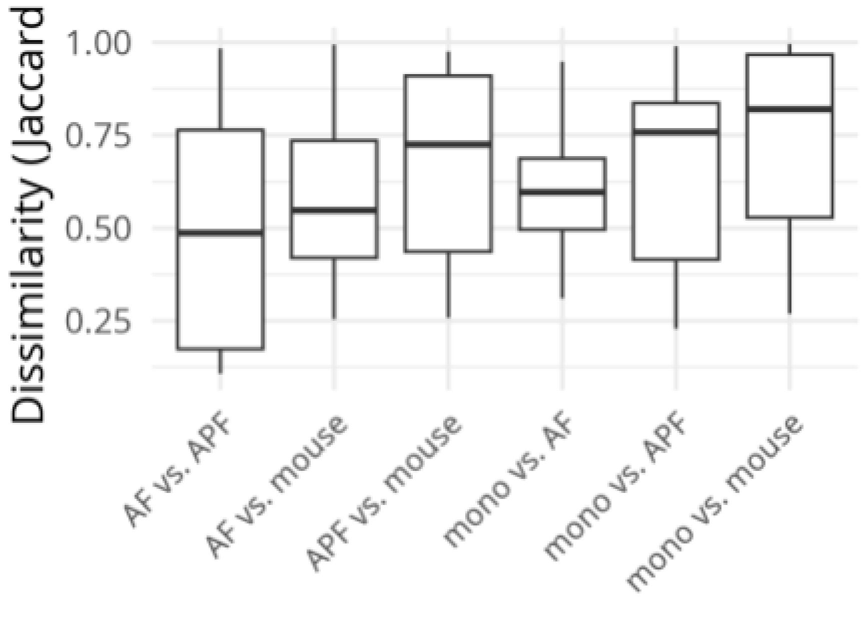


**Figure S16: Similarity of condition-specific metabolic models between conditions. Jaccard similarities of models based on model reactions are shown and compared pairwise between experimental conditions.**

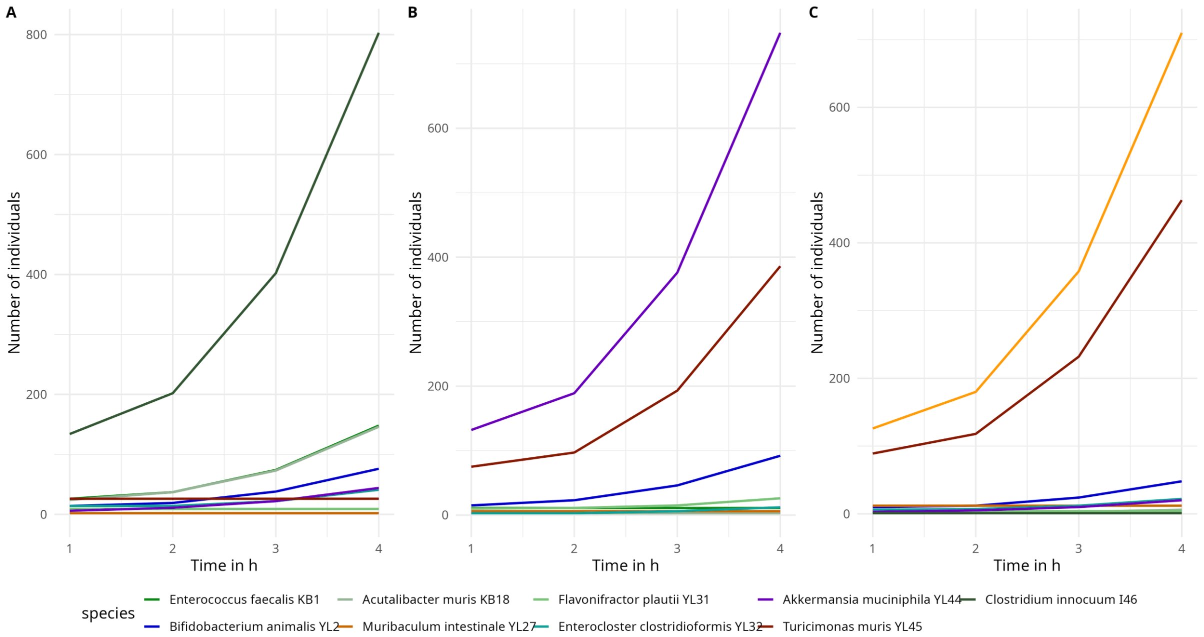


**Figure S17: Predicted growth curves of the OMM$^{12}$ community with AF (A), APF (B) medium and in mouse (C).**

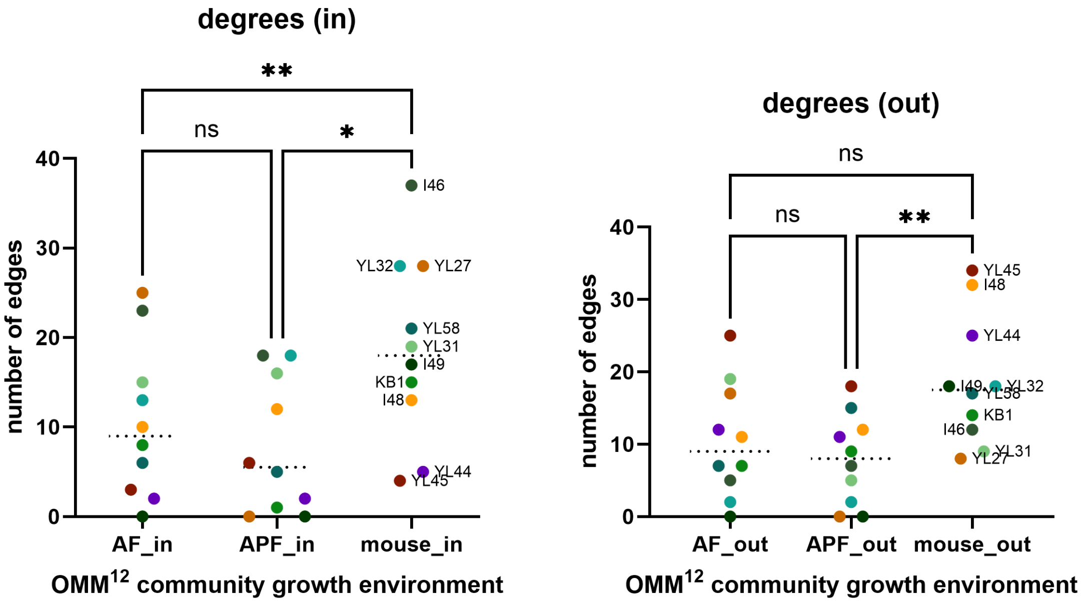


**Figure S18: Connectivity of species within the community network broken down into ingoing and outgoing edges.**

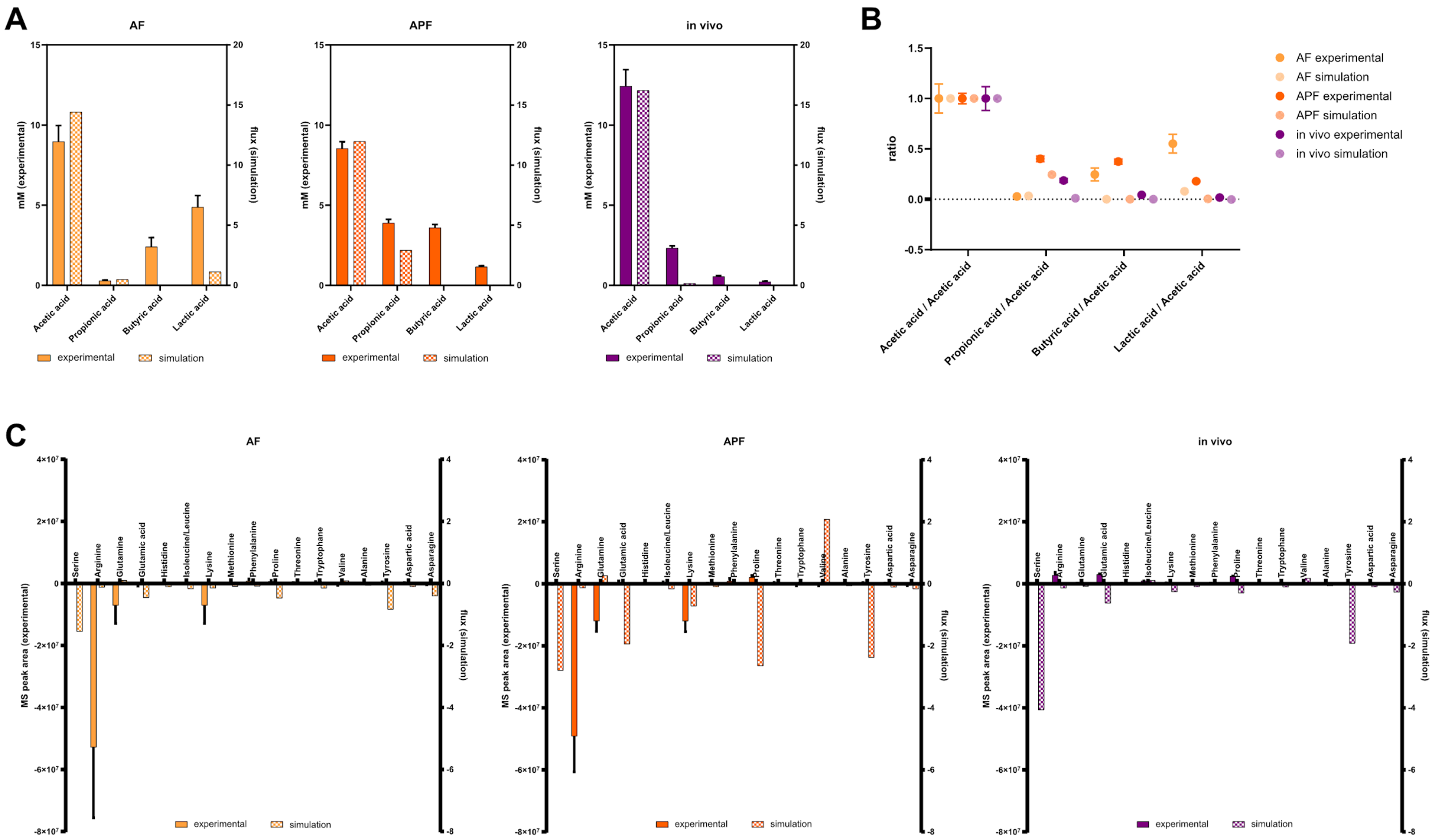

**Figure S19: Comparison of simulated short-chain fatty acid (A) and amino acid (C) production profiles as well as short-chain fatty acid ratios (B). Short-chain fatty acid concentrations are shown in mmol/l; amino acid signals as MS peak area. Experimental data are shown as mean +/- SEM.**

**Table S1: Cultivation times for proteomic samples**

| Species | Growth time after 10% inoculation | Volume |
|---|---|---|
| *E. faecalis* KB1 | 4h | 10ml |
| *B. animalis* YL2 | 5h | 15ml |
| *A. muris* KB18 | 48h | 50ml |
| *M. intestinale* YL27 | 14h | 15ml |
| *F. plautii* YL31 | 14h | 15ml |
| *E. clostridioformis* YL32 | 14h | 15ml |
| *A. muciniphila* YL44 | 24h | 15ml |
| *T. muris* YL45 | 48h | 50ml |
| *C. innocuum* I46 | 4h | 15ml |
| *B. caecimuris* I48 | 4h | 15ml |
| *L. reuteri* I49 | 5h | 20ml |
| *B. coccoides* YL58 | 5h | 15ml |
| OMM$^{12}$ | 24h | 15ml |

**Table S2: Data points marked in volcano plots (Figures 1D-H and S2)**
-> separate Excel file

**Table S3: full datasets for enrichment monoculture - community (Figures 1I-K, S3, S4)**
-> separate Excel file

**Table S4: full datasets for enrichment in vitro - in vivo (Figures 2E, S9, S10)**
-> separate Excel file

**Table S5: Full gene annotations of community expression data (Figures 2F and S7)**
-> separate Excel file

**Table S6: reference data for model validation**
-> separate Excel file

**Table S7: predicted exchange fluxes in the metabolic network models (Figure 4B)**
-> separate Excel file

**Table S8: reactions involving pyruvate in the I46 mouse CSM (Figure 4D)**
-> separate Excel file

**Table S9: expression of PTS system components in *C. innocuum* I46 and of the trehalose PTS system in *E. faecalis* KB1 (Figure 4**
-> separate Excel file

**Table S10: characteristics and identifiers for OMM[12] strains**

| Species | strain | DSM number | GenBank accession | RefSeq accession | Known substrates | Known products |
|---|---|---|---|---|---|---|
| *Enterococcus faecalis* | KB1 | DSM32036 | CP065317 | NZ_CP065317.1 | glucose, arginine, serine, trehalose | lactate, formate, ornithine |
| *Bifidobacterium animalis* | YL2 | DSM26074 | CP065311 | NZ_CP065311.1 | tryptophan, xylose, threonine, glucose | acetate, lactate |
| *Acutalibacter muris* | KB18 | DSM26090 | CP065321 | NZ_CP065321.1 | glucose | acetate |
| *Muribaculum intestinale* | YL27 | DSM28989 | CP065316 | NZ_CP065316.1 | glucose | propionate, butyrate, acetate, arginine isobutyrate, isovalerate, methylbutyrate |
| *Flavonifractor plautii* | YL31 | DSM26117 | CP065315 | NZ_CP065315.1 | lysine, glutamine, glucose | butyrate, propionate ,xanthine, acetate |
| *Enterocloster clostridioformis* | YL32 | DSM26114 | CP065314 | | cysteine, fucose, glucose | $H_2S$, butyrate, acetate |
| *Akkermansia muciniphila* | YL44 | DSM26127 | CP065322 | NZ_CP065322.1 | fucose | propionate, succinate |
| *Turicimonas muris* | YL45 | DSM26109 | CP065313 | NZ_CP065313.1 | Aspartate, asparagine | succinate, acetate, ammonia |
| *Clostridium innocuum* | I46 | DSM26113 | CP065320 | NZ_CP065320.1 | trehalose, glucose | butyrate, formate, valerate, caproate |
| *Bacteroides caecimuris* | I48 | DSM26085 | CP065319 | NZ_CP065319.1 | inulin, xylan, xylose, glucose | propionate, succinate, acetate, isobutyrate, isovalerate, methylbutyrate |
| *Limosilactobacillus reuteri* | I49 | DSM32035 | CP065318 | NZ_CP065318.1 | arginine | succinate, acetate, lactate |
| *Blautia coccoides* | YL58 | DSM26115 | CP065312 | NZ_CP065312.1 | $H_2$, alanine, trehalose, glucose | acetate, succinate |
